# Uncovering the heterogeneity of pancreatic endothelial cells using integrative and comparative single cell gene expression analysis

**DOI:** 10.1101/2023.04.19.537540

**Authors:** Safwat T. Khan, Neha Ahuja, Sonia Taib, Shabana Vohra, Ondine Cleaver, Sara S Nunes

## Abstract

The pancreatic islet vasculature displays tissue-specific physiological and functional adaptations that support rapid glucose sensing and insulin response by β-cells. To uncover the transcriptomic basis of this specialization, we performed a meta-analysis of multi-organ single cell RNA sequencing atlases employing a unique strategy to avoid transcriptomic contamination. We identified biologically relevant genes involved in sphingosine-1-phosphate-mediated insulin-secretion (*PLPP1, RDX, CDC42EP1*), islet basement membrane formation (*SPARC, COL15A1*), endothelial cell (EC) permeability (*PLVAP, EHD4*), membrane transporters (*CD320, SLCO2A1)* and developmental transcription factors (*NKX2-3, AHR)*. These were validated *in silico* in independent datasets. We further established the first integrated transcriptomic atlas of human pancreatic ECs and described two unique capillary subpopulations: exocrine and endocrine pancreas ECs. We validated the spatial localization of key markers using RNAscope™ and immunofluorescence staining on mouse pancreatic tissue cross-sections. Our findings provide novel insights into pancreatic EC heterogeneity and islet EC function with potential implications in therapeutic strategies.

## Introduction

Endothelial cells (ECs) are the interface for tissue-blood exchange of oxygen, nutrients, and metabolic waste products. ECs also secrete soluble factors, known as angiocrine factors, that are important for tissue morphogenesis, repair, and homeostasis(*1*). Furthermore, ECs adopt unique tissue-specific characteristics to meet the functional needs of the surrounding parenchyma such as in the brain(*2*) and liver(*3*). Similarly, the pancreatic Islets of Langerhans are made up of several hormone-secreting cell-types, such as insulin-secreting β-cells and glucagon-secreting α-cells(*4*), which directly regulate blood glucose levels and are thus intimately connected to the vasculature. In fact, despite making up only 1-2% of the pancreatic mass(*5*), islets are densely vascularized and receive up to 10-20% of the blood volume to the pancreas(*6–7*). Pancreatic ECs also display pore-like structures, where the endothelium tapers off into a thin diaphragm, called fenestrations, which increase the permeability of small molecules across the endothelium and facilitate the exchange of factors between the blood and extravascular tissue. Compared to exocrine vasculature, the islet vasculature is known to have ∼10 times the number of fenestrations(*5*). Importantly, an intricate cellular-cross talk between ECs and β-cells that is imperative for proper β-cell development, proliferation and insulin secretion has also been demonstrated *in vivo*(*8–9*). Furthermore, endothelial aberrations in the islets have been implicated in the pathophysiology of both type 1 and type 2 diabetes mellitus(*10–11*). However, the specific mechanism whereby islet fenestrations are formed, how islet ECs mediate insulin-glucose transport and their role in diabetes is not well understood.

The recent use of single cell/nucleus RNA sequencing (scRNAseq/snRNAseq) technologies has allowed major advances in the characterization of tissue-specific endothelial signatures for various organs (*12–13*). However, characterization of islet ECs is challenging. This is because the exocrine pancreas is primarily composed of acinar cells, which secrete high levels of hydrolytic enzymes. When the whole pancreas is dissociated into single cells/nuclei, these enzymes interfere with sequencing data. As a result, the majority of the pancreas datasets have sequenced isolated, *in vitro* cultured islets(*14–23*), which have low number of ECs sequenced. Therefore, these datasets do not allow for dissecting *intra* tissue pancreatic endothelial heterogeneity (exocrine vs endocrine). Further, (*24*) comparison of the gene expression of pancreas/islet ECs to other cell types in the pancreas does not allow for extracting information regarding endothelial specialization across tissues. Therefore, to dissect *inter* tissue endothelial heterogeneity, it is more relevant to compare pancreatic ECs to ECs in other tissues.

However, integrating EC data from numerous publicly available studies that have sequenced different organs is challenging. This is because any downstream analysis is highly susceptible to the sequencing library preparation technique(*24*) making it difficult to ascertain actual biological gene expression differences from that conjured by batch-effects. Therefore, for an unbiased comparison between different tissue EC sequencing data, they must be acquired using the same methodology. As such, we have looked towards multi-organ gene expression atlases, which sequenced and processed all tissues under a standardized workflow.

Kalucka et al.(*25*) have generated one such comprehensive murine endothelial gene expression atlas, unfortunately it does not contain pancreas ECs. Other studies(*26–27*) have also generated organotypic signatures for ECs by subsetting only ECs from the Tabula Muris atlas(*28*), which contains scRNAseq data on ∼ 100,000 adult mouse cells generated using the FACS-based SmartSeq2 assay. However, they identified acinar-specific genes as top pancreas-specific EC genes, including genes such as *CPA1, CLPS, CEL, CELA3B, CTRBB1* and *PNLIP*. This highlights an issue of RNA contamination, which likely results from the lysis of acinar cells during single cell dissociation.

Therefore, to define a biologically relevant pancreas EC enriched gene signature we have analyzed ECs subsetted from two different multi-organ atlases, the adult mouse Tabula Muris and the human fetal Descartes (*29*). We use these two multi-organ atlases to generate their respective EC-only atlases and then determined cross-atlas conserved candidates for a pancreas EC enriched gene signature. We validate these candidates *in silico* on a third EC atlas, obtained from the Tabula Sapiens(*30*). Further, to avoid misidentifying ambient contaminant genes as pancreas EC enriched genes, we curated a list of endothelial genes by analyzing four pancreas scRNAseq/snRNAseq datasets. We then use this list to cross-reference discovered gene lists with. Using our analysis pipeline, we identify genes conserved across three different multi-organ gene expression atlases that constitute a signature that is enriched in pancreas ECs compared to other tissue ECs. Top genes of interest identified are involved in processes coherent with islet endothelial functions, such as sphingosine-1-phosphate mediated insulin secretion (*PLPP1, RDX, CDC42EP1)*, islet basement formation (*SPARC, COL15A1)*, endothelial barrier integrity *(HEG1)*, vascular permeability *(PLVAP, EHD4)* and membrane transport *(CD320, SLCO2A1)*. In addition, we introduce the first integrated human pancreatic EC atlas derived from three human datasets and dissect intra-tissue pancreatic EC heterogeneity. Using this integrated atlas, we described and validated *in silico* and *in situ* the existence of transcriptomically distinct endocrine-specific and exocrine-specific capillary subpopulations. Our approach provides a comprehensive overview of the inter and intra-tissue transcriptomic landscape of pancreatic ECs. This will help us better understand the role of islet microvessels in regulating glucose homeostasis and advance research in diabetes therapy.

## Results

### Comparative analysis of multi-organ gene expression atlases to define a pancreas enriched gene signature for endothelial cells

To generate a transcriptomic profile of pancreatic ECs, we have conducted comparative analysis against ECs from other tissues using multi-organ gene expression atlases and integrated multiple human pancreatic EC sequencing data to dissect the inter and intra-tissue pancreatic endothelial heterogeneity respectively (Fig. 1). First, we defined a pancreatic EC-enriched gene signature, i.e. a set of endothelial genes whose collective expression pattern differentiates pancreatic ECs from that of other tissue ECs. To do so, we subsetted and pooled all cells previously annotated as endothelial in other studies, from all available organs from the adult mouse Tabula Muris and fetal human Descartes multi-organ single cell atlases (Table 1). This generated an EC atlas for the Descartes and Tabula Muris respectively (Fig. 2 A-F). We identified differentially expressed genes (DEGs) (log2FoldChange > 0.25) upregulated in pancreas ECs compared to ECs from all other tissues in each atlas (Fig. 2C and 2F). This generated a list of DEGs enriched in pancreas ECs from the Descartes and another for the Tabula Muris EC atlases (Table S1 and S2). However, closer inspection of these DEGs revealed a significant number of acinar and endocrine-specific genes, likely resulting from ambient RNA contamination from surrounding exocrine and endocrine cells (*31*). While there are tools that can tease out ambient RNA expression such as SoupX(*32*) and DecontX(*33*), their methodologies depend on detecting ambient RNA that is expressed in all cell types obtained from the same sample. However, when comparing ECs across tissues that were isolated and sequenced tissue-by-tissue, the ambient RNA contamination present in pancreatic ECs is different from that of other tissues. As such, the contaminating DEGs could be identified as *de novo* genes. Therefore, we elected to not use *SoupX* or *DecontX*.

**Figure 1.**
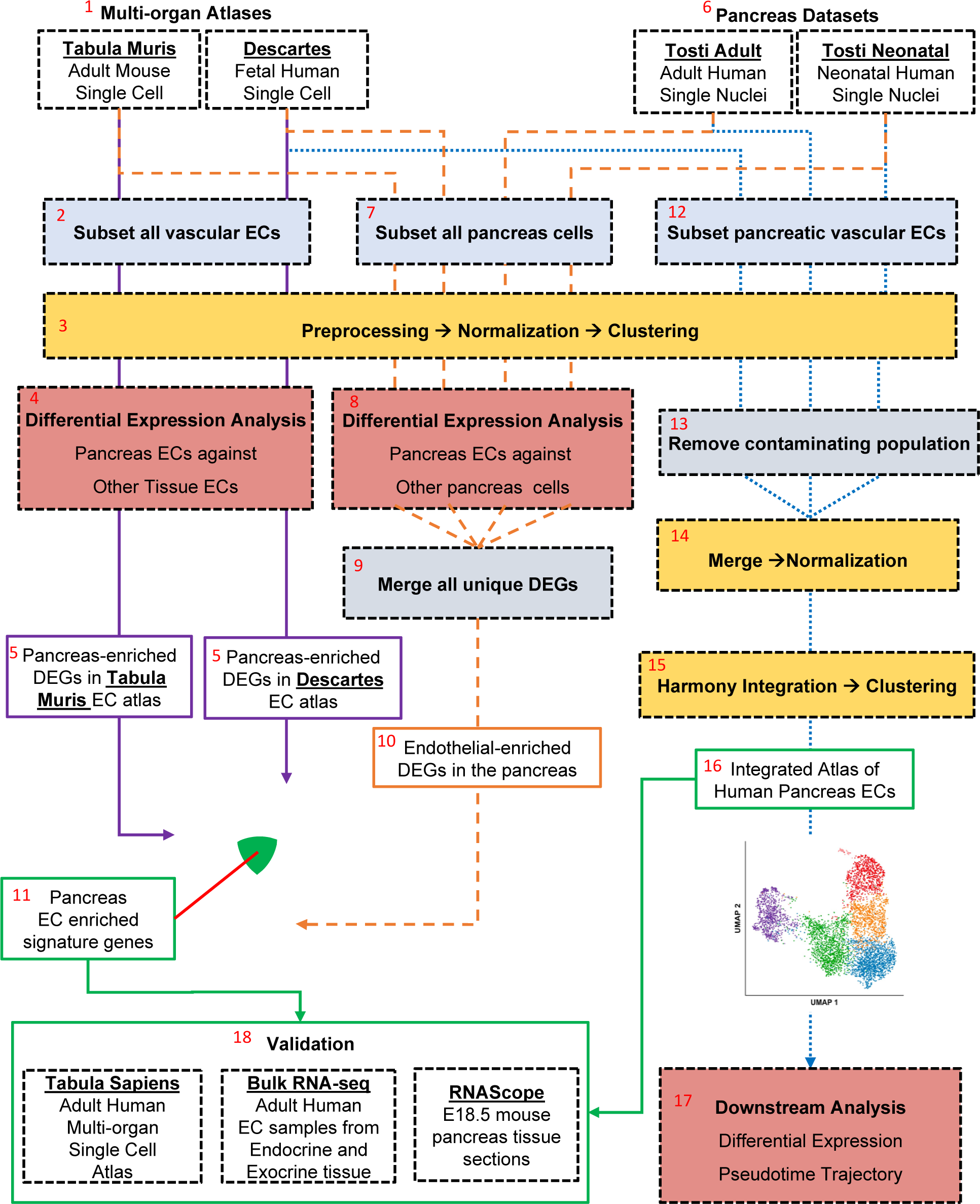
Analysis strategy to uncover a transcriptomic profile of pancreatic endothelial cells. This schematic summarizes the analysis conducted to obtain a biologically relevant pancreas enriched EC gene signature and a human pancreatic integrated atlas. Pancreas enriched EC signature genes lie at the overlap between pancreas-enriched differentially expressed genes (DEGs) derived from endothelial atlases and endothelial-enriched DEGS derived from pancreas datasets. Endothelial only atlases were derived from the human fetal Descartes and the adult mouse Tabula Muris multi-organ gene expression atlases (1) by subsetting and pooling all vascular ECs from different organs (2). After preprocessing, normalizing and clustering (3), DEGs are identified between pancreas ECs against other tissue ECs (4) for each atlas (5). Next, two pancreas only datasets from Tosti et al (6) and all pancreas cells were subsetted from the multi organ atlases (7). After preprocessing, normalizing and clustering these datasets, DEGs were identified between pancreas ECs and other pancreas cells (8). These DEGs were merged (9) to produce a list of endothelial-enriched DEGs derived from pancreas datasets (10). The overlap of this list and the pancreas-enriched DEGs from the EC atlases produces the pancreas-specific EC genes (11). Next, to create an integrated atlas of pancreatic ECs, all pancreatic vascular ECs were subsetted (12) from the human pancreas dataset sources (Descartes and Tosti). These ECs were individually processed, normalized and clustered. After which, contaminating clusters were removed (13). The remaining ECs were then merged and renormalized (14). The normalized counts were then integrated using Harmony (15), which produced the integrated pancreatic EC atlas (16). This atlas was used for further downstream analysis including DEG analysis within pancreatic EC subclusters and Pseudotime Trajectory analysis (17). Finally, the pancreas EC enriched gene signature (9) and populations of interest identified from the integrated atlas using Harmony (16) were validated *in silico* using the adult human Tabula Sapiens multi-organ gene expression atlas, bulk RNA-seq data of human exocrine and endocrine EC samples and in situ using RNAScope™ on embryonic day 18.5 (E18.5) mouse pancreatic tissue (18).

**Figure 2.**
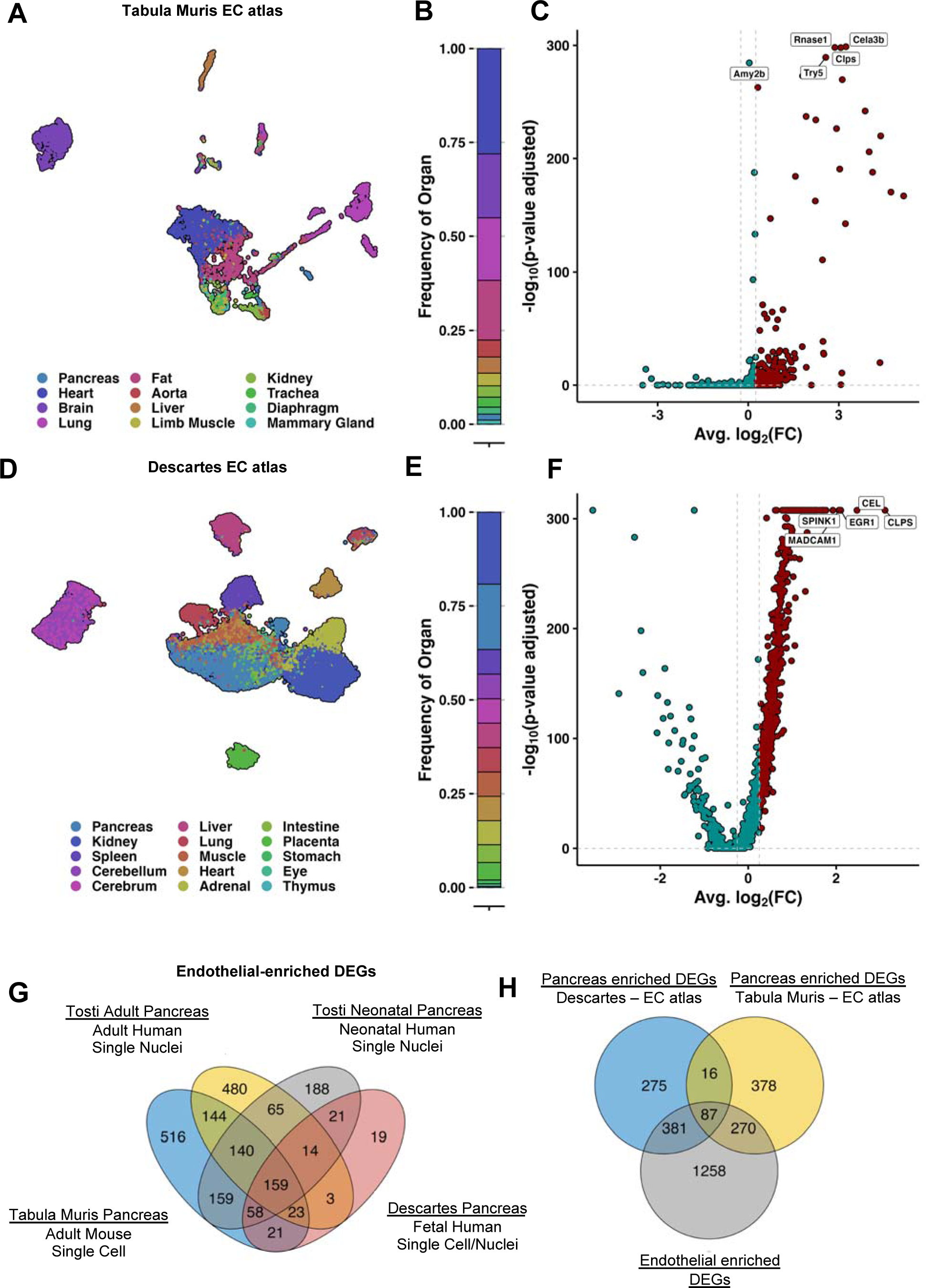
Determining a biologically relevant pancreas enriched endothelial cell (EC) gene signature. A-F) Shows the summary of endothelial atlases derived from the Tabula Muris (A-C) Descartes (D-F). A) and D) shows The UMAP visualization of single cell sequencing data for the ECs from all available organs. B and E) Highlight the tissue/organ ratios in the EC atlases, the colors correspond to the labelling in the preceding UMAPs. C and F) are volcano plots demonstrating differentially expressed genes (DEGs) in pancreas ECs against other organ/tissue ECs. Enriched DEGs (log2FoldChange > 0.25; Wilcoxon Rank Sum Test) are shown in red and the rest of the DEGs in blue. Further, the top 5 significant DEGs are labeled. G) Shows the overlap of significant endothelial marker DEGs (log2FoldChange > 0.25; Wilcoxon Rank Sum Test, Bonferroni-adjusted p-value < 0.05) derived from four different pancreas single cell/nuclei gene expression datasets (Tabula Muris, Descartes, adult and neonatal pancreas from Tosti et al.). Endothelial marker DEGs are defined as DEGs upregulated in vascular ECs in the pancreas compared to all other pancreas cell-types. Merging all unique DEGs from the different datasets produced the list of “endothelial-enriched DEGs”. H) Shows the overlap of pancreas-enriched DEGs (log2FoldChange > 0.25; Wilcoxon Rank Sum Test, Bonferroni-adjusted p-value < 0.05) derived from EC atlases obtained from the human fetal Descartes and adult mouse Tabula Muris gene expression atlas, and the previously derived merged list of endothelial-enriched DEGs. The overlap of all these sets produces the “pancreas enriched EC gene signature”.

**Table 1.** Summary of datasets used in this study.

| <u>Dataset</u> | <u>Citation</u> | <u>Assay</u> | <u>Sequencing Method</u> | <u>Age</u> |
| --- | --- | --- | --- | --- |
| Descartes<br>(Human) | Cao et al., Science, 2020(29) | Single Cell | Sci-RNA-seq3 | Fetal |
| Tabula Muris<br>(Mouse) | Tabula Muris Consortium (28) | Single Cell | SmartSeq2 | Adult |
| Tosti – Adult<br>(Human) | Tosti et al., Gastroenterology, 2021 (34) | Single Nuclei | 10X Genomics | Adult |
| Tosti – Neonatal<br>(Human) | Tosti et al., Gastroenterology, 2021 (34) | Single Nuclei | 10X Genomics | Neonatal |
| Tabula Sapiens<br>(Human) | Tabula Sapiens Consortium (30) | Single Cell | 10X Genomics | Adult |

Instead, to avoid selecting contaminant genes as part of the pancreas EC enriched gene signature, we cross-referenced the pancreas enriched DEGs identified from the DE analysis from the Descartes and Tabula Muris EC atlases, with a list of endothelial genes obtained by contrasting the pancreatic EC genes with genes from all other pancreatic cells. That was achieved by analyzing four different pancreas datasets: the scRNAseq pancreas data from the Descartes, Tabula Muris gene expression atlases, a human adult and a human neonatal snRNAseq pancreas datasets generated by Tosti et al.(*34*) (henceforth referred to as Tosti-Adult and Tosti-Neonatal, respectively) (Table 1, Fig. S1 and S2). For each dataset, we identified the significantly enriched DEGs (log2FoldChange > 0.25, Bonferroni-adjusted p-value < 0.05) enriched in the pancreatic vascular EC populations compared against all other non-vascular pancreatic populations, such as β-cells or acinar cells, excluding lymphatic ECs and perivascular cells, which could have overlapping gene expression with ECs (Table S3). We reasoned that any DEGs that are enriched in ECs cannot be contaminating genes from other populations, as the contaminating genes should have a higher expression in the cell-type it originates from. We observed limited overlap in the endothelial genes discovered in the four datasets. This is due to both biological and technical variability and highlights the ability of each dataset to detect unique endothelial genes in the pancreas that cannot be detected in the sample type or by the methodology used in the other datasets. As such, to ensure our list does not exclude potential endothelial genes, we combined all 1,996 unique DEGs identified from the four datasets into one list (Fig. 2G). We refer to these DEGs as the curated list of endothelial enriched DEGs.

Next, we cross referenced the pancreas enriched DEGs from the EC atlases (Descartes and Tabula Muris) with the curated list of endothelial-enriched DEGs (Fig. 2H) (Table S4 and S5), and any genes that did not overlap. Most of the top EC DEGs in both atlases are contaminating genes (Fig. 3A and B). Among the remaining DEGs, there were 87 genes which were common between all three lists i.e. enriched in pancreas ECs in the Descartes EC atlas, the Tabula Muris EC atlas and were all endothelial enriched genes (Fig S3). This indicates that these genes are highly conserved and potentially relevant, as they were identified as being endothelial enriched genes that are also pancreas enriched across two different EC atlases from different species (human and mouse), sequencing methodologies (10x and smartseq2) and age (adult and fetal). We refer to these 87 genes as candidate genes for a ‘pancreas EC enriched gene signature’ (Table 2). Hierarchical clustering of the different organ ECs based on the expression of these candidate signature DEGs allowed pancreas ECs to cluster separately, in its own branch, in both the Descartes and Tabula Muris EC atlas (Fig. 3C and D).

**Figure 3.**
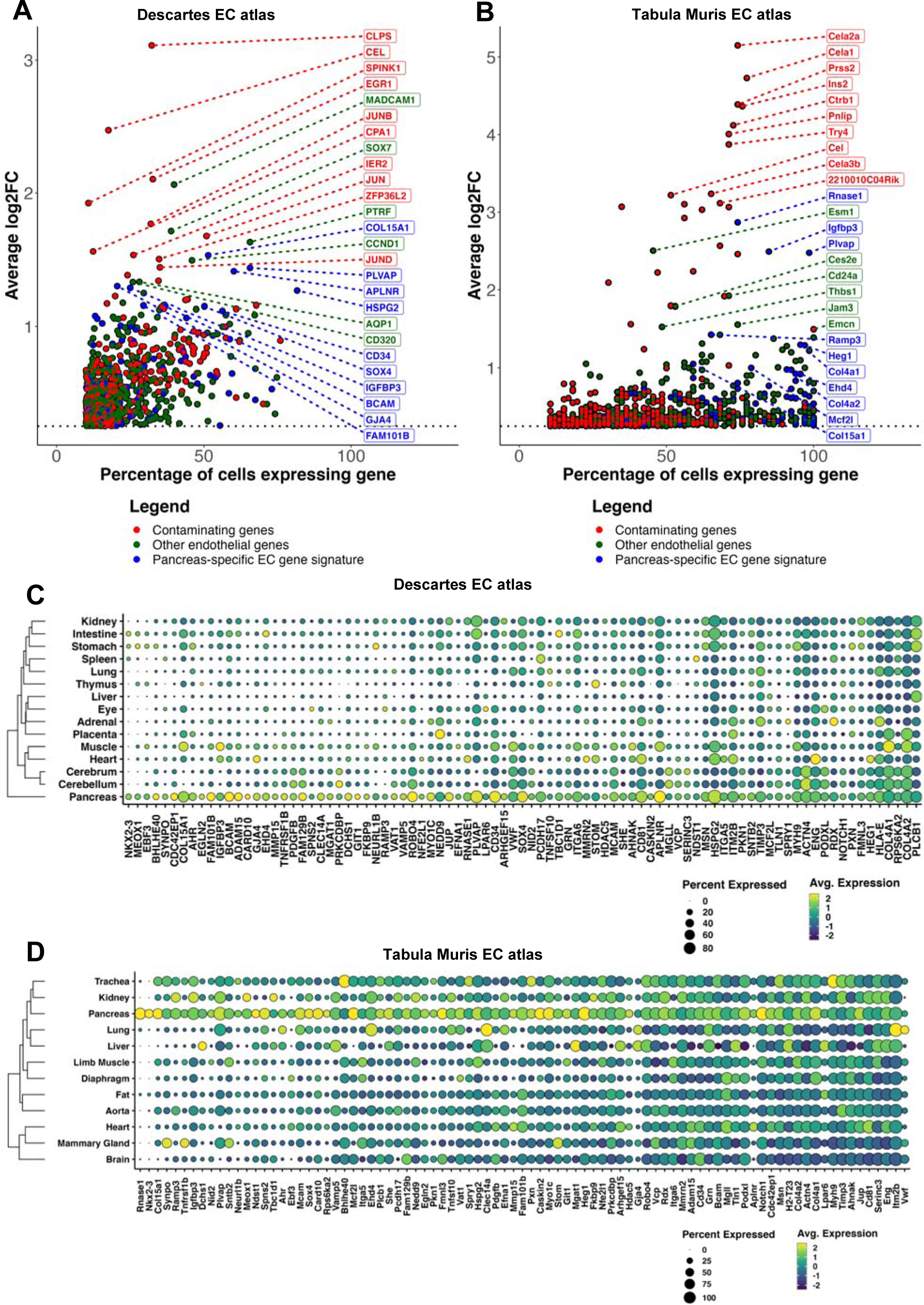
Assessment of the pancreas-enriched DEGs in the Descartes and Tabula Muris endothelial cell (EC) atlases. A) and B) demonstrate a scatterplot for all significant (log2FoldChange > 0.25; Wilcoxon Rank Sum Test, Bonferroni-adjusted p-value < 0.05) pancreas-enriched DEGs identified from the Descartes and Tabula Muris EC atlases. The y-axis is average log2FC of gene expression in the pancreas ECs vs other tissue ECs and the x-axis is percentage of ECs expressing the gene. Potential contaminating genes, are colored in red, pancreas EC enriched signature genes are in blue and the rest of the endothelial-enriched genes are in green. C) and D) demonstrates the individual average scaled log-normalized expression of the pancreas EC enriched signature genes in the Descartes and Tabula Muris EC gene expression atlas, respectively. Genes are ordered in decreasing order based on the fold-change in percentage of ECs expressing gene in the pancreas against other tissues/organs. Pancreas-specificity of this signature is demonstrated using hierarchical clustering of genes based on Euclidian distances between the average scaled expression of the DEGs across the different tissues/organs.

**Table 2.**
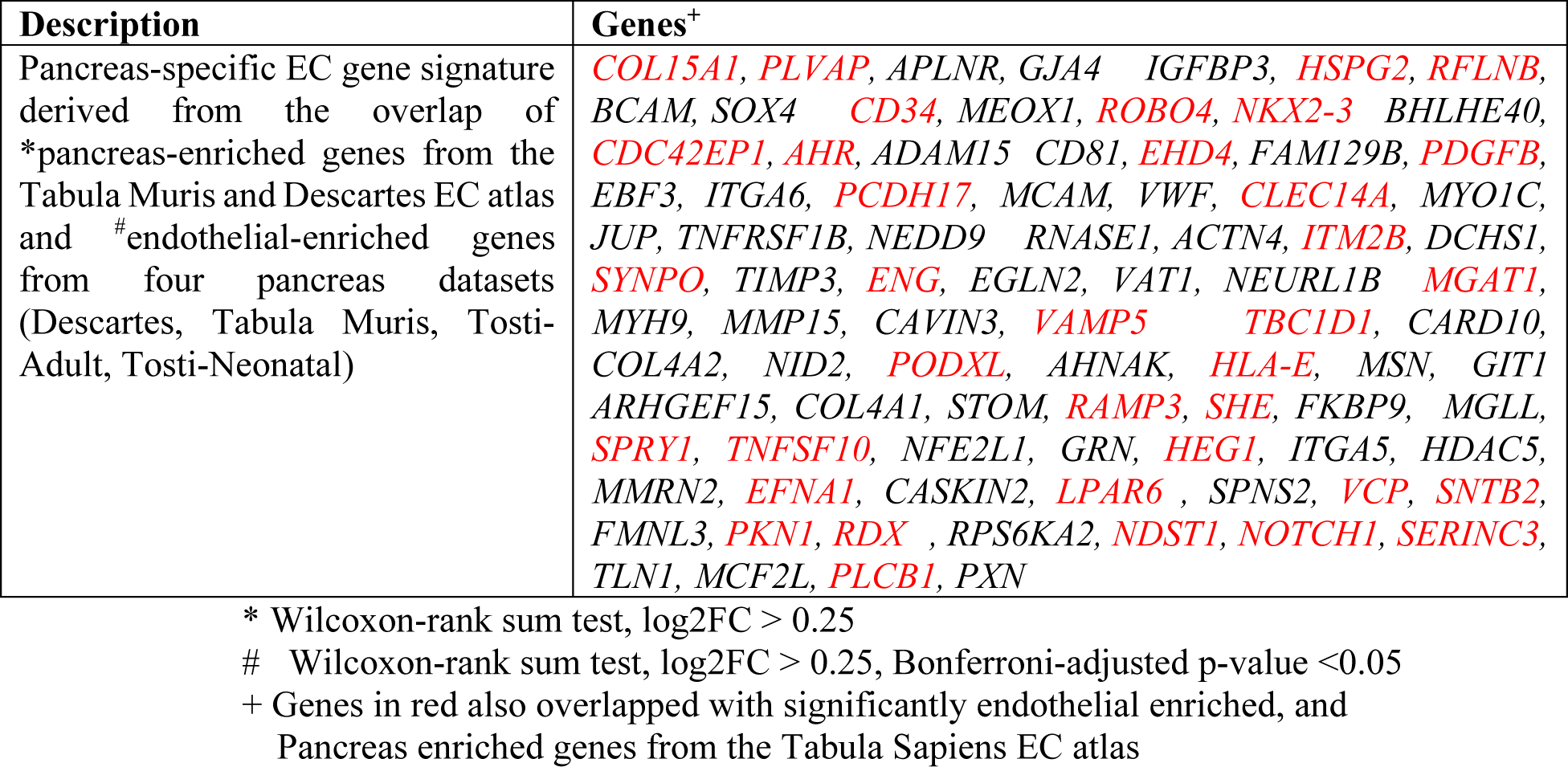
Relevant gene lists.

### Validation of pancreas EC enriched gene signature in an independent multi-organ atlas

To further increase confidence in the specificity of the pancreatic EC gene signature, we assessed its expression in an independent, third multi-organ atlas, not used to derive this signature. For this purpose, we analyzed the recently published adult human Tabula Sapiens gene expression atlas. Its corresponding EC atlas was generated by subsetting all annotated ECs (Fig. 4A-B), as done for other atlases above. We then assessed the collective expression of all 87 candidate genes using Seurat’s module score(*35*), which employs the scoring strategy described by Tirosh, et al(*36*). We showed that the module score for the candidate genes was highest in the EC population in the pancreas, compared to other EC populations in all other tissues (Fig. 4C). This corroborated that the 87 genes constitute a pancreatic EC enriched gene signature. Hierarchical clustering based on average scaled expression of the candidate signature DEGs, also clustered pancreas ECs separately from all other organ ECs (Fig. 4D). We next assessed which DEGs were the top scoring genes that drove the pancreas-specificity in the Tabula Sapiens EC atlas.

**Figure 4.**
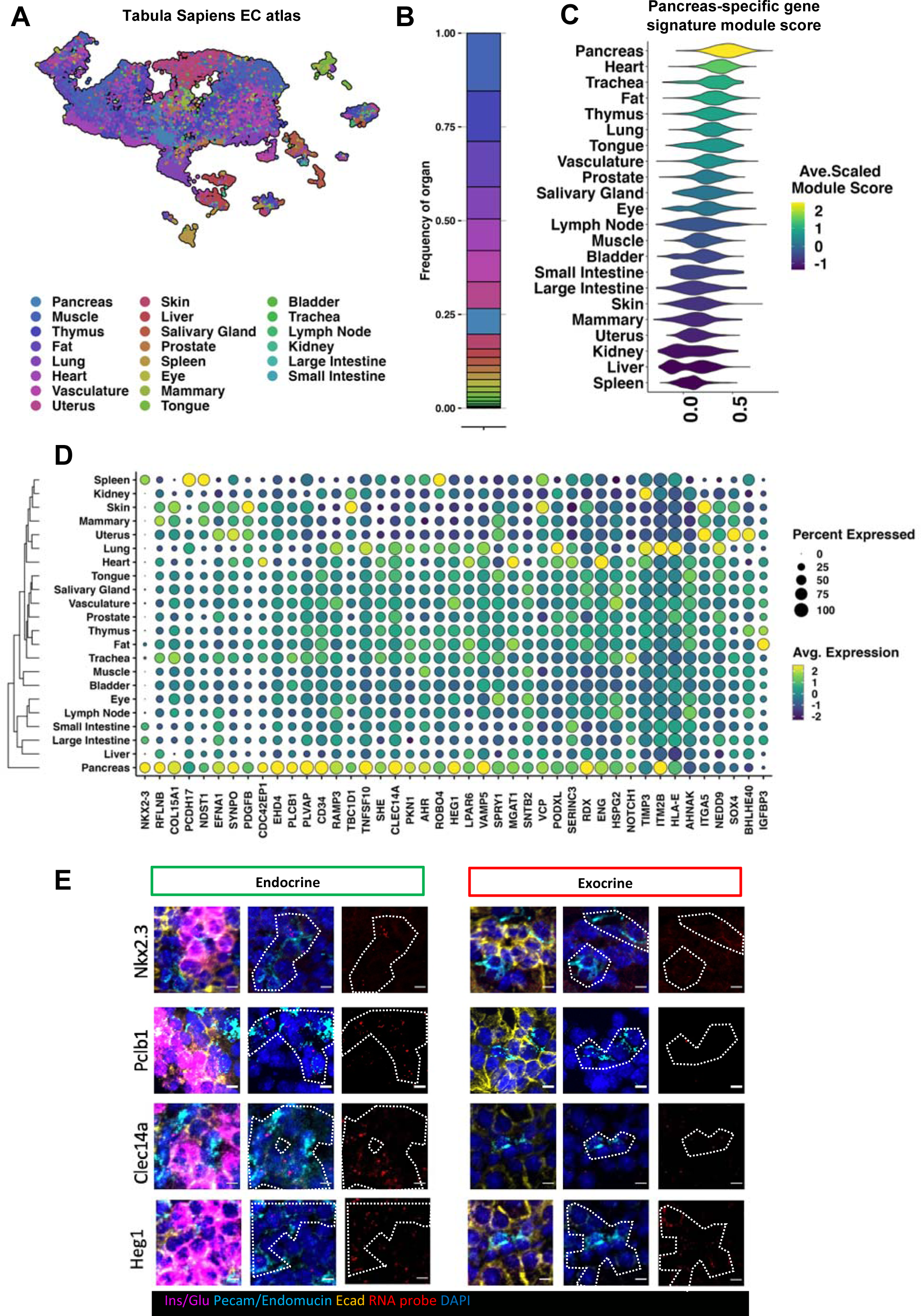
Validating the pancreas enriched endothelial cell (EC) gene signature in a third independent endothelial atlas obtained from Tabula Sapiens. A) shows a UMAP of all organ endothelial cells (EC) in the Tabula Sapiens atlas. B) Shows the ratios of each organ/tissue in the EC atlas and is colored corresponding to the preceding UMAP legend. C) demonstrates the module scores derived by assessing the collective expression of the pancreas EC enriched gene signature using Seurat’s AddModuleScore function in the different EC populations of the Tabula Sapiens EC atlas. Each violin is colored based on the average scaled module score for that cluster. D) demonstrates the individual log-normalized average scaled expression of the pancreas EC enriched signature genes in the Tabula Sapiens EC atlas. Hierarchical clustering based on Euclidian distances between the average scaled expression of the DEGs across the different tissues demonstrates specificity to the pancreas. E) demonstrates *in situ* images of E18.5 mouse pancreas tissue bound by RNAscope™probes (red) for selected pancreas EC enriched signature genes. Endocrine and exocrine regions are identified by immunofluorescence (IF) co-staining of insulin and glucagon (pink), which stains cells of the islets; vasculature is identified by co-staining of pecam-1 and endomucin (neon); all other epithelial cells are stained by e-cadherin (yellow); and the nuclei are stained by DAPI (blue). The white dotted outline demonstrates manually outlined regions rich in pecam1-endomucin co-staining.

Of these DEGs, *NKX2-3* was one of the top enriched genes in pancreatic ECs compared to other tissue ECs (Fig 4D). This gene was also one of the top pancreas-enriched genes in both the Descartes and Tabula Muris EC atlases (Fig. 3C and D). Other signature genes of interest validated in the Tabula Sapiens include *PKN1, HEG1*, *PLVAP*, *EHD4*, *COL15A1*, *CLEC14A* and *PLCB1*. We have further confirmed the vascular expression of *Nkx2-3* and other highly enriched genes (*Plcb1, Clec14a, Heg1*) *in situ* in E18.5 mouse pancreas (Fig. 4E). Importantly, among the signature genes validated to be pancreas enriched in the Tabula Sapiens EC atlas, we also observe several genes potentially involved in S1P-mediated regulation of endothelial barrier integrity via cytoskeletal alterations regulated by the Rho GTPase signaling pathway such as *AHR* (*37*)*, CDC42EP1*(*38*)*, RDX* (*41*), *SYNPO*(*39*)*, RFNLB* (or *FAM101B*). Conversely, *MSN* (*41*) and the *SPNS2* (*40*) were also present in the signature genes but were not validated in the Tabula Sapiens EC atlas. We also assessed other genes involved in S1P signaling such as sphingosine kinases, phospholipid lipases and S1P receptors in the Tabula Sapiens (Fig S4). We observed that pancreatic ECs had the highest expression of *S1PR1* and *PLPP1*(*41*) compared to other pancreatic cell-types, including β-cells, which did not express any of these genes directly. *PLPP1* was also the most enriched in pancreas ECs compared to other tissue ECs. We further identified all pancreatic EC-enriched genes in the Tabula Sapiens EC atlas. Similar to the other EC atlases, the top pancreas enriched DEGs were contaminating genes (Fig. S5A, Table S6). As before, we excluded contaminating genes by cross-referencing these DEGs with the curated list of endothelial enriched DEGs. In total, among all the pancreas enriched DEGs from the Tabula Sapiens EC atlas, 366 DEGs were also endothelial enriched (Fig. S5B). We then compared these with the endothelial and pancreas enriched DEGs from the Descartes and Tabula Muris EC atlases. Overall, 36 endothelial and pancreas enriched DEGs overlapped across all 3 atlases (Table 2, Fig. S5C and D).

In addition, 51 candidate genes overlapped between the Descartes and Tabula Muris EC atlases but were not enriched in the Tabula Sapiens EC atlas. There were also genes that only overlapped between the human Descartes and Tabula Sapiens EC atlases, but not the mouse Tabula Muris EC atlas. This may constitute human-specific pancreatic EC enriched genes. In total, 105 genes uniquely overlapped between pancreas enriched DEGs from the Descartes EC atlas and the Tabula Sapiens EC atlas (Fig. S5C-D). The top 10 DEGs by log2FC in pancreas ECs compared to ECs from other in the Tabula Sapiens EC atlas include: *CD320*, *SLCO2A1*, *IGFBP4, TMEM88, TXNIP, HLA-C*, *SPARCL1, SOX18* and *FAM13C*. 59 DEGs also uniquely overlapped between the two adult atlases, Tabula Sapiens EC atlas and Tabula Muris EC atlas, potentially identifying adult specific pancreas EC genes (Fig. S5C and D). The top 10 DEGs by log2FC from this list in the Tabula Sapiens EC atlas include; *TGFB2R, PLPP1*, *ITGA1, GAS6, GPR146, CCN2, ATOH8, KDR* and *FAM167B*. The remaining 166 DEGs that are unique to Tabula Sapiens are also of interest (Table S6).

### Defining the intra-tissue endothelial heterogeneity in the pancreas

Next, we assessed the intra-tissue EC heterogeneity in the pancreas. To maximize our transcriptomic resolution, we used Harmony(*42*) to integrate the pancreatic ECs across the 3 human datasets: the fetal pancreas EC data from Descartes (Descartes-Fetal), Tosti-Adult and Tosti-Neonatal (Fig. 5A). Read count matrices for ECs from each dataset were preprocessed, log-normalized and clustered as described in the methods section. No batch correction method was applied to the read counts to specifically account for differences in the source (Descartes or Tosti). This approach of only conducting log normalization on raw counts followed by wilcoxon rank sum test DE analysis has been shown to outperform more sophisticated batch correction tools (*43*). Unsupervised clustering of the pancreatic EC datasets revealed several contaminating clusters within each pancreatic EC dataset, which did not express canonical endothelial genes (*CDH5, KDR, VWF*) or co-expressed non-endothelial contaminating genes, such as acinar and endocrine genes. To ensure these populations did not interfere with proper integration of pancreatic ECs across datasets, the contaminating clusters were removed prior to integration. As a result, 3,857 ECs, 1,023 ECs and 576 ECs were used for integration from the Descartes-Fetal, Tosti-Adult and Tosti-Neonatal datasets, respectively.

**Figure 5.**
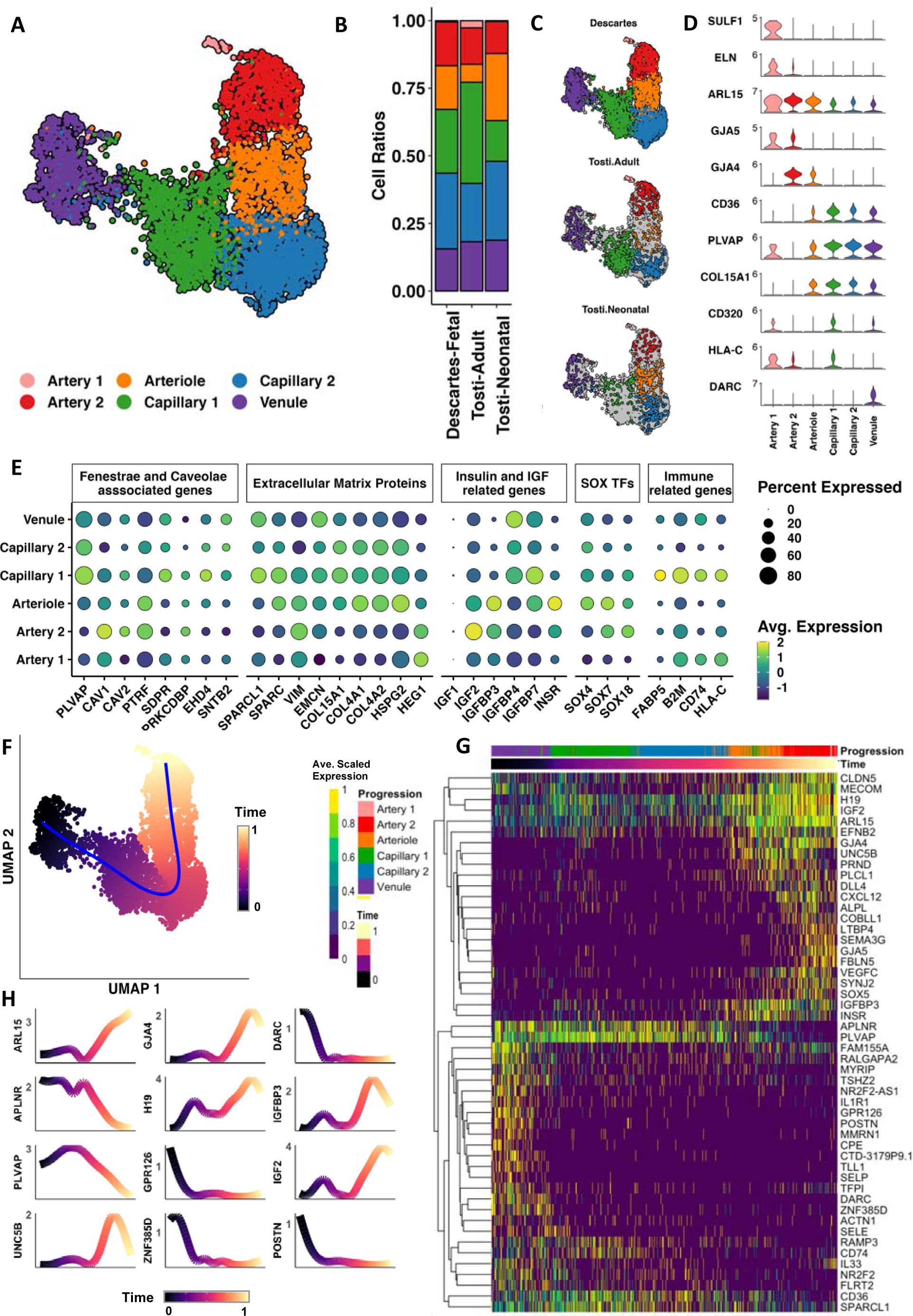
Integrated map of human pancreatic endothelial cells (ECs) Three different human pancreatic EC datasets were derived from their parent data and integrated using Harmony. This includes the fetal human pancreas dataset from the Descartes gene expression atlas and the neonatal and adult pancreas dataset from Tosti et al. A) shows the resulting integrated UMAP using the harmony transformed principal component values. B) Shows the percentage distribution of different subpopulations in each dataset colored according to the labelling of the preceding UMAP. C) shows the UMAP of the integrated map split by dataset (Descartes-Fetal, Tosti-Adult, Tosti-Neonatal). D) demonstrates violin plots showing expression of markers used to annotate different clusters in the integrated atlas. F) shows the UMAP of the integrated map colored by pseudotime inferred through trajectory analysis using SCORPIUS. The trajectory is colored in blue. G) shows the heatmap demonstrating the change in average scaled expression of the top 50 unique markers determined by Wilcoxon Rank Sum Test for each subcluster across pseudotime. Genes are hierarchically clustered based on the Euclidian distance between the average scaled expression of the genes. H) Shows the expression of the top 12 genes across pseudotime ranked by importance to trajectory interference using SCORPIUS.

This produced an atlas of human pancreatic ECs with 6 different subpopulations. These unique subpopulations were present in approximately equal proportions across the individual datasets (Fig. 5B and C). Based on gene expression patterns, we discovered that these populations corresponded to the topographical location of ECs along the vascular tree (Fig. 5D). Three arterial populations were identified by the expression of *GJA5*(*44*) *GJA4* (*45*) and *EFNB2* (*46*). Artery 1 population was additionally enriched for elastin (*ELN*) and *SULF1*, which are involved in artery extracellular matrix (ECM) formation(*47*). The arteriole population co-expressed arterial markers and *CD36*, which has also been shown to be differentially expressed in capillary ECs in the lung(*12*). *CD36* was also highly expressed in the two capillary populations, capillary 1 and capillary 2, alongside *PLVAP* and *COL15A1*. We also annotated one venule population based on the expression of the venule marker *DARC* (*48*).

DE analysis (Table S7) between the subpopulations revealed genes involved in caveolae and fenestrae formation, ECM production, insulin-like growth factor (IGF) signaling, SOXF transcription factors, immune-activity, and membrane transport. Therefore, we assessed other genes in these families to identify potential trends that correspond to vascular hierarchy (Fig. 5E). Notably, compared to capillary 2, capillary 1 had higher expression of several caveolae and fenestrae associated genes such as *PLVAP*(*49*), *SDPR* (or *CAVIN2*)(*50*) and *EHD4*(*51*). It also had higher expression of several immune-related genes, including *FABP5*(*52*) and *CD74*(*53*), *ITM2A, B2M, HLA-C, HLA-B*. Higher immunological activity and presence of fenestrae are characteristics that have previously been attributed to intra-islet endocrine capillaries in comparison to exocrine capillaries(*54*). This suggests that capillary 1 and capillary 2 subpopulations are islet and exocrine capillary ECs, respectively. Other genes highly enriched in capillary 1 included *SPARC, SPARCL1*, *IGFBP7*, *SLCO2A1, CD320* and *SLC2A3*. Importantly, many of these genes overlapped with the pancreatic EC enriched genes identified above (Table 2, Table S4 and S5). By contrast, capillary 2 displayed a more generic expression pattern with no significantly enriched genes above the log2FC > 0.25 threshold compared to capillary 1. This could be due to the lack of statistical power in the integrated atlas or due to lack of read depth in the datasets, as we did identify numerous non-significant (Bonferroni adjusted p-value > 0.05) DEGs or significant DEGs with log2FC < 0.25 in capillary 2. The arteriole population also had an interesting gene expression pattern, with high expression of *HEG1*, *SOX7*, *IGFBP3* and *INSR*.

Pseudotime trajectory inference of the integrated pancreatic EC map on the UMAP space using SCORPIUS, a machine learning tool which has been benchmarked to be one of the best trajectory methods based on overall performance(*55*) (Fig. 5F), predicted a trajectory from the venule population to the artery populations, through the capillary populations. This further supports capillary annotation. Gene expression of top markers (Table S7) for each vascular bed subpopulation across pseudotime also demonstrated a continuous gene expression pattern across the vascular topography rather than discrete endothelial subpopulations (Fig. 5G). We also identified genes that predict the order of the cells along pseudotime, ranked by an importance score derived using a Random Forests algorithm employed by SCORPIUS (Fig. 5H). This allowed us to identify the expression pattern of both established (*GJA4, GJA5, DARC*) and under studied arterio-venous markers for differentiating ECs along the vascular tree. Of these genes, we found that capillaries have peak expression of *PLVAP* and *APLNR*. Another gene of interest is *UNC5B*, which shows peak expression in the arteriole ECs and has been recently shown to regulate blood-brain barrier integrity through interaction with netrin-1(*56*). Several other genes (*POSTN, GPR126* and *ZNF385D)* match the expression pattern of *DARC*, which peaks specifically in venule ECs, suggesting these could potentially be novel marker genes for venous ECs.

### Validation of an islet-specific EC cell population and signature

To confirm that the capillary 1 EC sub-population corresponded to the islet ECs, we re-analyzed a bulk RNA sequencing dataset (*57*) of isolated ECs from endocrine (islets) and exocrine pancreas from 8 adult human donors. Principal component analysis demonstrated that the endocrine sample for donor #8 is an outlier (Fig. S6), hence donor #8 endocrine and exocrine samples were removed before downstream analysis. We then conducted unsupervised hierarchical clustering of the remaining endocrine and exocrine samples of the scaled log-transformed counts per million (CPM) values using the DEGs that were identified in the integrated atlas between capillary 1 and capillary 2 (Fig. 6A). This enabled the separation of the exocrine and endocrine samples in separate branches, thus further corroborating that the DEGs in the integrated atlas define exocrine and endocrine ECs. We also assessed whether the pancreas EC enriched signature that we established is enriched in the islet capillary EC population in the bulk RNAseq data. When we assessed the collective module score of the signature genes, we found that the endocrine EC samples were indeed enriched for this score (Fig. 6B). DE analysis between the endocrine and exocrine bulk RNAseq samples using DESeq2 (Table S8) demonstrated that 23 of the 87 signature genes in the pancreas EC enriched signature were also significantly (log2FC > 0.5, adjusted p-value <0.05) different between endocrine and exocrine EC samples (Fig. 6C). 19 out of the 23 were significantly enriched in endocrine ECs including *PLVAP, NKX2-3, COL4A1, COL4A2,* and 4 were significantly enriched in exocrine samples; *MEOX1, PCDH17, NEURL1B* and *TNFSF10* . This is again consistent with the integrated atlas, as we see that *NEURL1B* and *PCDH17*, had higher expression in capillary 2 in the integrated atlas, while *MEOX1* and *TNFSF10* expression were higher in other subpopulations than capillary 1 and capillary 2 (Fig. 6D). This also suggests that the exocrine tissue harbors other ECs subtypes than just capillaries. We also assessed the presence of other genes of interest shown by us to be enriched in capillary 1 compared to capillary 2 in the integrated atlas in the bulk RNAseq data. Of those, *SPARC* was significantly enriched in the endocrine samples according to the DESeq2 analysis (Fig 6C). *CD320, SLCO2A1, EHD4, IGFBP7* also had higher expression in endocrine samples but were not significantly enriched (adjusted p-value >0.05) (Fig S7).

**Figure 6.**
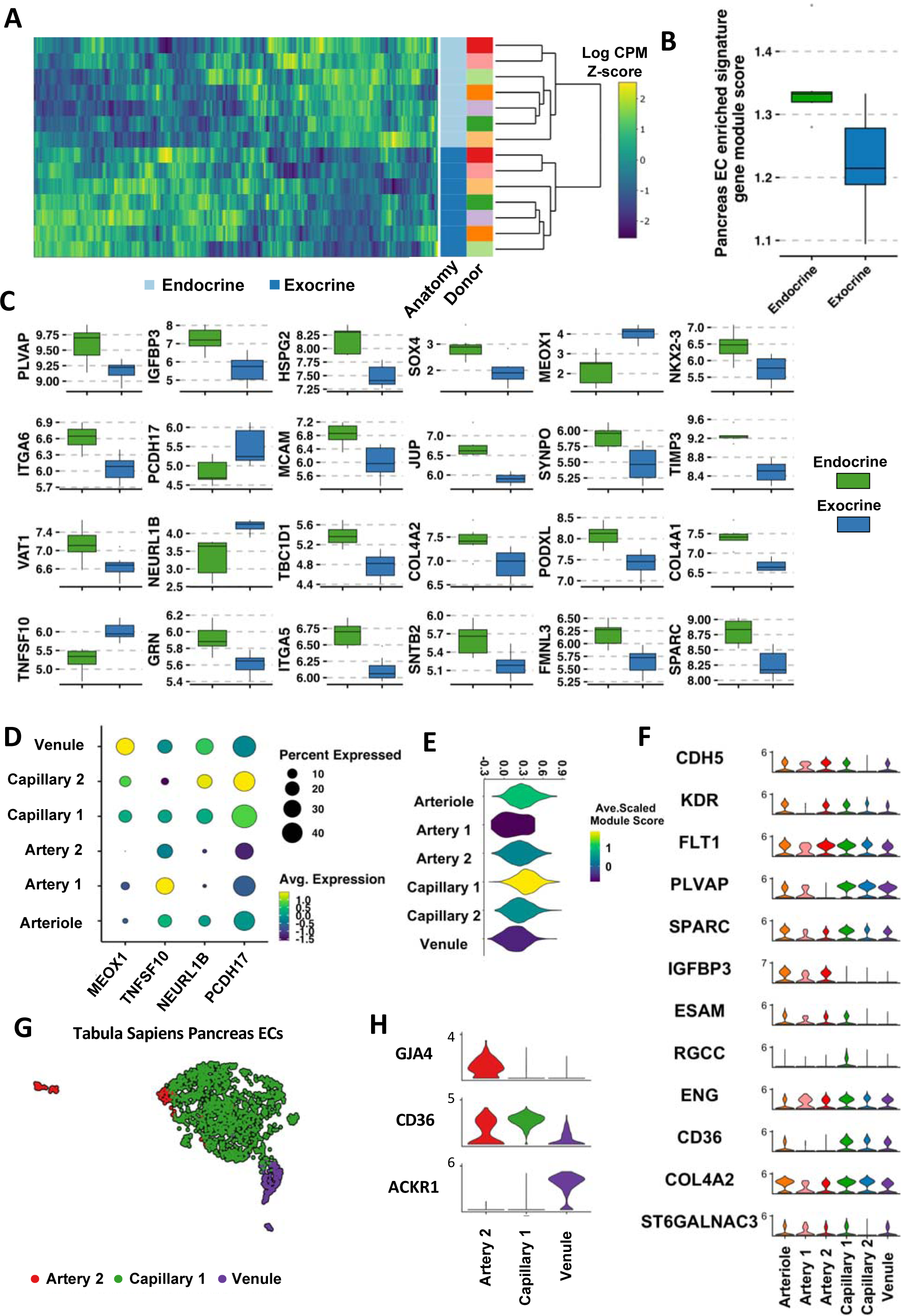
In silico validation of an islet capillary subpopoulation and specificity of pancreas EC enriched gene signature in islet capillary subpopulation. A) Shows the heatmap of log counts per million (CPM) z-scores for DEGs identified between capillary 1 and capillary 2 in the integrated atlas in the bulk RNA-seq endocrine and exocrine EC samples sequenced by Jonsson *et al*.. Hierarchical clustering based on Euclidian distance of the samples based on the log counts per million (CPM) z-scores allows clustering of samples based on anatomy. B) Shows a box plot of the collective module score for the pancreas EC enriched signature determined using Seurat’s AddModuleScore function for the bulk RNA-seq exocrine and endocrine samples. C) Shows box plots for all pancreas EC enriched signature genes and SPARC that were also determined to be significantly different (log2FC > 0.5, adjusted p-value <0.05) using DESeq2 between the exocrine and endocrine EC samples. D) Shows the expression in the integrated pancreas EC atlas of the 4 pancreas EC enriched signature candidate genes, which were found to be significantly enriched in exocrine EC samples in the bulk RNA seq data according to the DESeq2 analysis. E) Demonstrate expression of canonical markers identified in literature that are specific to endothelial cells (ECs) in islet single cell datasets in the integrated pancreatic EC atlas. F) Demonstrates the module scores derived by assessing the collective expression of the pancreas EC enriched gene signature using Seurat’s AddModuleScore function in the different EC subpopulations in the integrated EC atlas. G) Shows the Tabula Sapiens pancreatic ECs, annotated with Seurat’s reference mapping using the integrated pancreatic EC atlas as the reference, in order to validate presence of vascular bed subpopulations. D) Shows the expression of the markers identified in the integrated pancreatic EC atlas for the subpopulations predicted in the Tabula Sapiens pancreatic ECs.

Similar to the bulk RNAseq data, (*21–23*)among the subpopulations identified in the integrated atlas, the capillary 1 subpopulation had the highest enrichment for the module score for the pancreas EC enriched signature (Fig 6E), followed by the arteriole population. This suggests that it is the capillary 1 subpopulation that is driving this signature in pancreatic ECs and that the arteriole population is also specialized. This further supports that capillary 1 is the more specialized endocrine/islet-specific capillary population, while capillary 2 is the more generic exocrine capillary population. Recent publications(*21–23*) have also conducted scRNAseq of isolated islet cells and have uncovered markers for the endothelial population, which should majorly constitute the islet capillary ECs as well the feeding arteriole ECs. We’ve assessed these markers in the integrated map and demonstrated that capillary 1 highly expressed all these islet endothelial markers except *IGFBP3*, which as we demonstrated, is enriched in the arteriole population (Fig. 6F). This confirms the annotation of capillary 1 as islet capillary ECs. It is to be noted however, that these papers identified markers by comparing the ECs to non-ECs, as such not all genes are unique to pancreas ECs compared to other tissue ECs (Fig S8). We also applied a cell-type annotation transfer mapping method on the Tabula Sapiens pancreas EC dataset, using the integrated pancreatic EC atlas as the reference (Fig. 6G). Three different subpopulations were identified in the Tabula Sapiens pancreas: artery 2, capillary 1 and the venule subpopulations. These annotations were corroborated by assessing the expression of marker genes *GJA4, CD36* and *DARC* (or *ACKR1*) in these clusters (Fig. 6H). The majority of the cells were annotated as capillary 1, with some cells mapping to the artery 2 and venule population. No capillary 2 ECs were annotated, suggesting that ECs were likely damaged during the enzymatic digestion process in Tabula Sapiens. However, it could also be a caveat of the annotation transfer method, which was made for annotation of different cell-types and may not be able to tease out the differences between the EC subpopulations at the current resolution and read depth.

Finally, to validate *in situ* that the transcriptomic signature for capillary 1 and capillary 2 in the integrated map correspond to EC populations in the endocrine (or intra-islet) and exocrine capillaries respectively, *in vivo*, we assessed the expression of 3 DEGs of interest, *PLVAP, SPARC*, and *NKX2-3*, which were enriched in capillary 1 in the integrated atlas (Fig. 7A) and that was enriched in endocrine ECs against exocrine ECs in the bulk RNA seq data. We conducted RNAscope™ and IF staining of these genes *in situ* on E18.5 mouse pancreatic tissue (Fig. 7B). RNAscope™ data demonstrated that a substantial portion of the probes, 70.1% (± 3.44 %) for *Sparc* and 49.6% (± 5.44 %) for *Nkx2-3,* were co-localized within the vessels (Fig. 7C). This is consistent with our analysis that demonstrate *NKX2-3* is also expressed by perivascular cells (Fig. S3). IF staining and quantification of *Plvap* showed that it had over two times higher fluorescence intensity (3970AU ± 571) in endocrine *Pecam1/Endomucin*^+^ capillaries compared to exocrine capillaries (1513 AU ± 268) (Fig. 7D). Further, *Sparc* in the endocrine vessels had over 3 times as much RNA expression level as measured by area of bound probe (0.0416 µm² ± 0.0086) compared to the exocrine vessels (0.01234 µm² ± 0.0012) (Fig. 7D). RNAscope™ of *Nkx2-3* did not show any statistically significant difference in endocrine vs exocrine vessels, however, the mean area of probe binding was higher within the islets. Overall, the *in situ* expression pattern of endocrine and exocrine capillary ECs, further corroborates our capillary annotation for the integrated pancreatic EC map.

**Figure 7.**
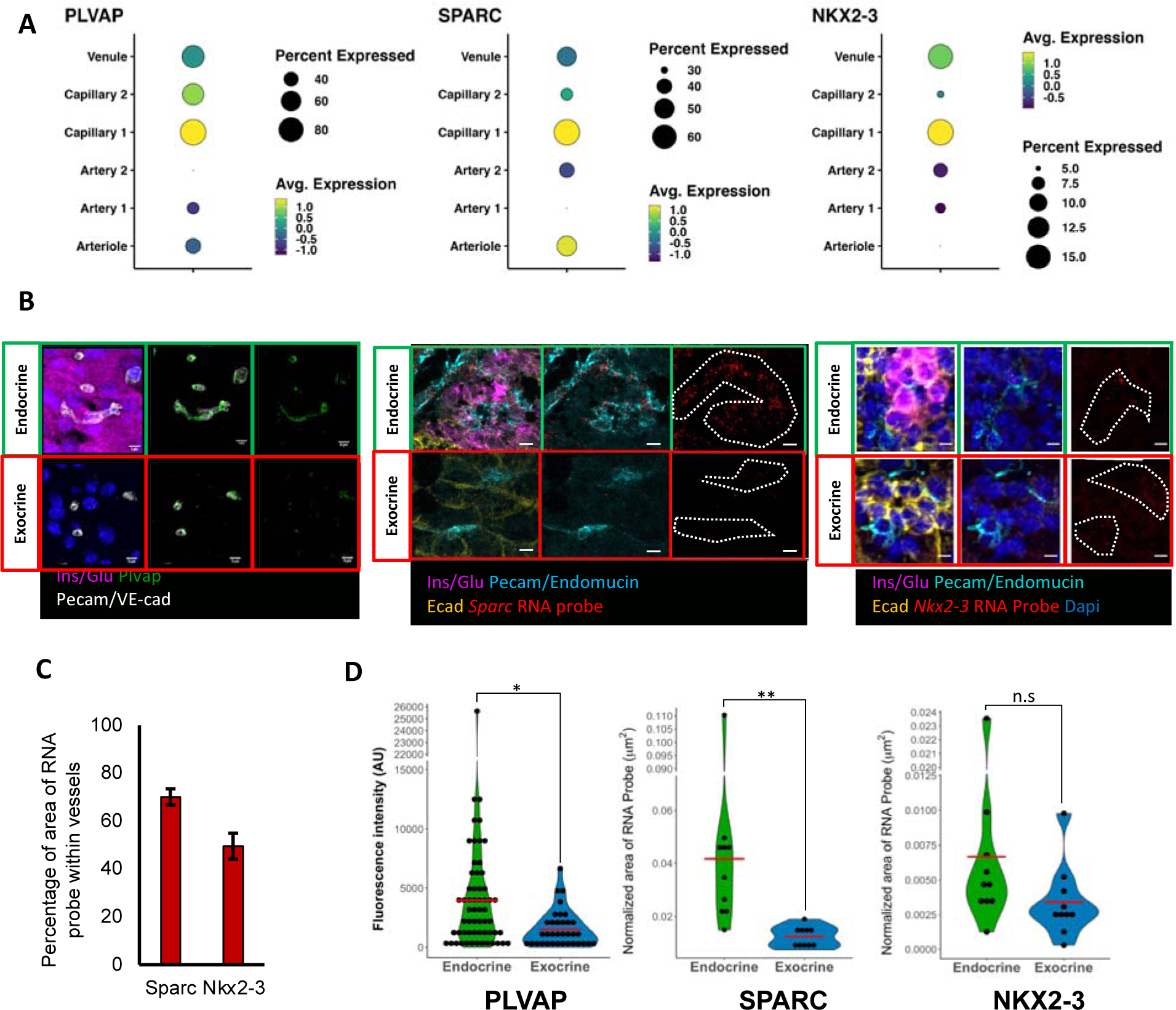
*In situ* validation of islet-specific EC capillary population. A) demonstrates the expression of 3 different DEGs that were enriched in the capillary 1 population of the integrated EC atlas. These genes include *PLVAP* (left)*, SPARC* (middle) and *NKX2-3* (right). B-D) demonstrate *in situ* validation of the 3 selected capillary 1 DEGs on E18.5 mouse pancreatic tissue cross-sections using RNAScope™ and Immunofluorescence (IF) staining. B) demonstrates representative images taken of microvessels in exocrine and endocrine regions for IF staining (green) of Plvap (left) and RNAscope™probe staining (red) for *Sparc* (middle) and *Nkx2-3* (right) in tissue cross-sections. Pink shows IF co-stain of glucagon and insulin, which highlights the islets from the exocrine tissue for all images; green for Plvap IF and red for *Sparc* and *Nkx2-3* RNA probes. Images highlight microvessels co-stained with Pecam1 and endomucin (cyan); yellow shows E-cadherin staining. The scale bar represents 5 μm. C) shows the average percentage of probe bound within microvessels outlined by a blinded researcher based on only Pecam1/endomucin IF staining across all tissue cross-sections. D) Demonstrates quantification of fluorescence intensity in arbitrary units (AU) for Plvap and RNAscope™area of probe binding (μm^2^) for *Sparc* and *Nkx2-3* in endocrine and exocrine microvessels, where each vessel was considered 1N. One-tailed t-test statistics was used to calculate significance (* - p-value <0.05, ** - p-value <0.01, *** - p-value <0.001).

## Discussion

Endothelial cells demonstrate a remarkable extent of intra and inter-tissue heterogeneity in structure, function, and molecular characteristics. Omics technologies now enable us to dissect this organotypic heterogeneity down to the single cell resolution. Here, we have analyzed different multi-organ gene expression atlases and have established a unique, cross-species conserved, *in silico* and *in situ* validated pancreatic EC-enriched gene signature. This signature included genes involved in insulin-secretion, fenestrae and caveolae formation, basement membrane components, endothelial barrier integrity, vascular permeability, and key developmental TFs. Furthermore, we have established the first integrated atlas of human pancreatic ECs using three recently published human pancreatic EC datasets from fetal, adult, and neonatal human samples. This is similar to previously generated integrated atlases for ECs in the brain(*13*) and lung(*12*) and allowed us to identify distinct subpopulations with unique gene expression patterns corresponding to the topographical location of ECs along the vascular tree. Specifically, we describe two unique capillary subpopulations. We then validated that these capillary subpopulations indeed correspond to endocrine (islet) and exocrine capillary ECs by assessing their gene expression patterns in bulk RNAseq data from isolated islet EC and exocrine ECs from adult human donors. We also validated the expression of select species-conserved markers *in situ* on E18.5 mouse pancreatic tissue using RNAscope and IF staining.

The importance of profiling the transcriptomics of pancreatic ECs is clear when considering the integral nature of ECs in β-cell development and function, and its relevance in diabetes. The pancreas-specific EC gene signature can shed insight into the development of specialization in glucose homeostasis and insulin transport within the islet microvasculature. Further, owing to the tissue-specific roles of ECs, there is a recent push towards generating induced pluripotent-stem cell (iPSC)-derived organotypic ECs for tissue-specific vascular regeneration, which remain outstanding challenges in regenerative medicine, and to build physiologically relevant 3D vascularized tissue models *in vitro*(*58–59*).

Past attempts at generating a transcriptomic signature of pancreatic ECs have inadvertently introduced bias from contaminating/ambient RNA from lysed acinar cells (*26–27*), which skewed the differential expression analysis. To circumvent this issue and uncover a contaminant-free, gene signature for pancreatic ECs, we have cross-referenced the pancreas enriched EC genes with a list of endothelial enriched markers in the pancreas, i.e., they are expressed in ECs and not by possible contaminating populations such as endocrine and exocrine cells. It is worth mentioning that two different endothelial gene expression atlases (Descartes and Tabula Muris EC atlas) were used to derive the candidates for the pancreatic EC enriched signature, which was then *in silico* validated on a third, independent gene expression atlas, the Tabula Sapiens EC atlas. This signature was also specifically enriched in islet capillary ECs in both the integrated pancreatic EC atlas and the bulk RNAseq data. We also validated gene expression *in situ*, confirming the expression of several of the signature genes within the ECs of the pancreas. Altogether, this increased the confidence that the observed gene expression patterns are representative of the pancreatic microvasculature.

Specifically, we identified and validated *PLVAP*, which encodes for plasmalemma vesicle-associated protein that is present in the diaphragms of endothelial fenestrae and in caveolae (*49*), as being highly enriched in pancreatic ECs. This is true even when comparing to other fenestrated ECs such as of the kidney. *PLVAP* was also enriched in capillary 1 ECs compared to all other EC populations in the integrated atlas, including capillary 2. Consistent with this, PLVAP protein expression was validated *in situ* to be significantly higher in endocrine capillaries compared to exocrine capillaries. This indicated that capillary 1 was the endocrine population and capillary 2 was the exocrine population. In the integrated atlas, *SDPR* (or *CAVIN2*) and *EHD4* were also enriched in capillary 1 ECs compared to all other EC populations. *EHD4* was also identified in the pancreas-EC enriched gene signature. *SDPR* directly regulates caveolae formation and morphology (*50*) and *EHD4* colocalizes on the surface of caveolae(*51*). Altogether, this data shows higher concentration of caveolae and fenestrae in endocrine pancreatic ECs compared to exocrine ECs as well as other tissue ECs. This is functionally consistent with previous reports implicating caveolae in insulin and glucose transport(*60–62*).

Importantly, one of the top genes of interest across all three EC atlases was *NKX2-3* – which we validated *in situ* and showed that it was indeed expressed in the EC in the pancreas. *NKX2-3* also had higher expression in endocrine capillary ECs compared to exocrine capillary ECs in the integrated pancreas EC atlas and the bulk RNAseq dataset. However, we were unable to confirm that *NKX2-3* was enriched in endocrine capillaries using *in situ* hybridization. This could be due to the fact that only a fraction of ECs in the capillary 1 population expresses high levels of *NKX2-3*, making it difficult to tease out differences *in situ* between exocrine and endocrine vessels.

*NKX2-3* is a member of the NK2 family of homeobox-domain containing TFs. These are critical organ-specific developmental TFs(*63–64*). *NKX2-3* in particular is involved in the development of the spleen and gut(*65*), both of which arise from the endoderm, which also gives rise to the pancreas. Specifically, *NKX2-3* has been shown to be integral for proper vascular specification in these organs(*66–67*). Primarily, *NKX2-3* is involved in regulating the expression of mucosal vascular addressing cell adhesion molecule 1 (MAdCAM-1)(*98*), a tissue-specific adhesion molecule that facilitates the homing of leukocytes in vascular segments involved in immune surveillance and inflammation, such as the high endothelial venules found in lymph nodes and Peyer’s patches of the small intestine(*68–69*). Interestingly, MAdCAM-1 expression was highly enriched in pancreatic ECs in the fetal human Descartes EC atlas as well as the adult human Tabula Sapiens EC atlas, albeit very few pancreatic ECs expressed MADCAM1 in the Tabula Sapiens. Beyond MAdCAM-1, *NKX2-3* has also been shown to be involved in regulating expression of other important angiocrine factors such as endothelin-1, VEGF-A, adrenomedullin and endothelial nitric oxide synthase(*70–71*). The murine endothelial atlas described that *Nkx2*-3 was a tissue-specific marker for small intestine, colon and spleen ECs(*25*). Here, we established that *NKX2-3* was enriched in pancreatic ECs at even higher levels than these organs across all three EC atlases, and specifically in the islet capillary ECs compared to exocrine capillary ECs. This makes the *NKX2-3* TF an exciting target to study. Specifically, its role in pancreatic vascular specification should be further investigated.

We also described the presence of numerous genes enriched in pancreatic ECs that are related to S1P signalling. S1P is a bioactive lipid, which is secreted by platelets, red blood cells and ECs(*71*), that has garnered much research interest due to its multifaceted role in embryonic development, organ homeostasis and disease pathogenesis(*72*). In the pancreas, vascular derived S1P is involved in pancreas development(*73*). Further evidence suggests that extracellular S1P may induce glucose-stimulated insulin secretion in β-cells(*74–75*). In fact, exogenous S1P has been shown to improve β-cell proliferation in a type 2 diabetic mouse model(*76*). In this study, we highlight several genes involved in S1P signalling. This includes *AHR* a TF involved in vascular development(*77*) and angiogenesis(*78*) which has also been implicated in regulating S1P bioavailability(*37–79*); *CDC42EP1,* which binds to Cdc42 a member of the Rho family of GTPases, which in turn is involved in *S1P* signaling; (*37–79*)*RDX* (Radixin) and *MSN* (Moesin), which are members of the Ezrin-Radixin-Moesin (ERM)-actin binding complex and are integral in S1P-mediated endothelial barrier permeability(*80*); *RFLNB* (or *FAM101B*) and *SYNPO*, which are cytoskeletal genes consistent with S1P-mediated cytoskeletal alteration (*38*); *SPNS2*, which is a S1P transporter(*40*); and *PLPP1,* which dephosphorylates S1P (*41*). Taken together, these findings suggest that pancreatic ECs play a critical role in the production, transport, and metabolism of S1P. Notably, S1P receptor genes (S1PR1-5) were not detected in β-cells across all pancreas datasets, whereas pancreas ECs highly expressed *S1PR1* compared to other pancreatic cell-types. This suggests that S1P signaling primarily involves the islet ECs, which then propagates its effects to β-cells through other means to regulate their function. Further research is needed to fully elucidate the role of S1P and its associated proteins in the β-cell/EC axis.

Another pancreas EC enriched gene of interest is *SPARC*, also known as osteonectin, which is a member of the SPARC-family of matricellular proteins. Unlike conventional ECM proteins, matricellular proteins are non-scaffolding components of the ECM that are involved in a myriad of diverse and complex biological functions(*81*) such as regulating ECM remodelling, cell-matrix interactions and growth-factor activity. Importantly, it has been demonstrated *in vivo* that SPARC may regulate β-cell growth(*82*) and insulin secretion(*83*) and is essential for proper glucose homeostasis in a type 2 diabetic mouse model(*84*). Further, *SPARC* was found to be differentially expressed in the islets and in the plasma of diabetic patients(*85–86*). Here, we demonstrated that *SPARC*, and another member of the family, SPARC-like protein 1 (*SPARCL1),* also known as Hevin, are enriched in pancreatic ECs in the Tabula Sapiens EC atlas. *SPARC* and *SPARCL1* expression were also enriched in pancreas vascular ECs compared to all other pancreas cell-types, although stromal cells also expressed both genes as previously reported(*82*). Further, in the integrated pancreatic EC atlas as well as the bulk RNAseq data, we observed enrichment of *SPARC* in islet capillary ECs compared to the exocrine capillary ECs. *In situ* validation confirmed that islet capillaries had significantly higher *Sparc* expression.

Further, according to the integrated atlas, islet capillary ECs had higher expression of the vitamin B12 transporter *CD320*, prostaglandin transporter, *SLCO2A1* and glucose transporter *SLC2A3* (or *GLUT3*). *CD320* was also enriched in pancreas ECs in both human atlases. Therefore, the role of vitamin B12 and prostaglandin should be further investigated in pancreatic ECs. As these genes encode for membrane associated proteins, they could also act as potential biomarkers for islet capillary ECs along with the caveolae and fenestrae associated genes *PLVAP, EHD4* and *SDPR*.

Finally, we identified an arteriole population enriched for *IGFBP3* and *INSR* compared to the other EC populations in the integrated atlas. This indicates potentially important roles in insulin-signalling in the arterioles in the pancreas and is in line with a previously reported role for insulin signaling in promoting vasodilation via nitric oxide(*87*). Recently, it was established that pericytes can also regulate blood flow in islet capillaries in response to changes in blood glucose levels(*88*). Therefore, a mechanism whereby pre-capillary arterioles are regulating blood flow along with pericytes through interstitial insulin signaling from β-cells is likely.

Our study has limitations that need to be acknowledged. To relate genes of interest identified in this study to notable biological pathways, we examined the top genes and then consulted the literature to determine relevant processes. A more comprehensive approach would be to use enrichment analysis with a curated pathway database specific for ECs. However, current databases for gene set and pathway enrichment analysis such as REACTOME, KEGG and Gene Ontology have limited processes involved in vascular functions. Therefore, gene set enrichment using these databases may overlook important biological information present in the data. We also used Harmony to integrate pancreatic EC data from fetal, adult, and neonatal datasets. Harmony corrects the principal components obtained from dimensionality reduction techniques. It does not make any changes to the underlying raw count matrix of gene expression values. Therefore, it is not feasible to find DEGs between fetal, neonatal, and adult pancreatic EC datasets, as we cannot be sure whether the differences are due to batch effects. However, we can still compare gene expression across different populations found in all three datasets. Developing a normalization or transformation method that removes batch-effects in scRNAseq/snRNAseq gene expression is an ongoing area of intensive research (*89–90*). Finally, we did not conduct *in situ* validation on adult pancreatic tissue. This is because of the high levels of hydrolytic enzymes in the exocrine pancreas, which we found caused both tissue and RNA probe degradation, as well as autofluorescence, leading to nonspecific background staining that interfered with the accuracy and sensitivity of the staining.

Despite these limitations, we have succeeded in tuning out the transcriptomic noise that is inherent in pancreatic endothelial scRNAseq/snRNAseq data and determined a relevant and validated portrayal of the vascular transcriptomic architecture in the pancreas. Genes of interest and transcriptomic phenomena highlighted in this study reveal important functional roles of pancreatic ECs in glucose homeostasis and diabetes pathophysiology and identify important TFs that can aid future endeavours at generating pancreatic islet ECs for application in regenerative medicine and *in vitro* models.

## Methods

### Data Acquisition

Raw counts matrices and meta data for the scRNA-seq/snRNA-seq datasets used in this paper were derived from their respective online databases/repositories. For the Descartes gene expression atlas (https://descartes.brotmanbaty.org/), sparse gene (row) by cell (column) matrices along with cell annotation were downloaded for pancreatic and endothelial cells. For the Tabula Muris gene expression atlas, processed Seurat Robjects for all FACS-sequenced organ datasets were downloaded and the raw count matrices and meta data available for the cells were extracted (https://figshare.com/projects/Tabula_Muris_Transcriptomic_characterization_of_20_organs_and_tissues_from_Mus_musculus_at_single_cell_resolution/27733). Similarly, for the adult and neonatal pancreas data from Tosti et al., processed Seurat Robjects were downloaded (http://singlecell.charite.de/cellbrowser/pancreas/) and the raw count matrices and available meta data were extracted from the objects. For the Tabula Sapiens atlas, processed Seurat Objects containing all tissue endothelial cell data and pancreas cell data were downloaded and (https://figshare.com/articles/dataset/Tabula_Sapiens_release_1_0/14267219) the raw count matrices and available meta data were pulled from the objects . Bulk RNA seq counts were obtained from (https://www.ncbi.nlm.nih.gov/geo/query/acc.cgi?acc=GSE157546)

### Single cell analysis pipeline

Datasets were analyzed using Seurat (v4.0.1). Low quality cells with less than 200 unique genes and genes with 0 counts across all cells were removed. Cells with high unique feature counts (potential doublets or multiplets) above a certain threshold were omitted. The thresholds were determined for each dataset individually based on distribution of feature counts. Cells with high mitochondrial content were filtered out. Data was normalized using log normalization and highly variable genes were detected using variance-stabilizing transformation (vst). The normalized data was centered and scaled, and cell cycle scores were generated for each cell using lists of genes expressed in S Phase and G2M Phase. Scoring followed the strategy outlined by Tirosh et at. This data was re-scaled with regression of cell cycle scores and percentage of mitochondrial genes using the vars.to.regress parameter. Dimensionality reduction was performed using the “RunPCA” function, the top 30 principal components were used for non-linear dimension reduction using Uniform Manifold Approximation and Projection (UMAP), with the RunUMAP function. The first 10 principal components were used for Seurat’s default graph-based clustering using the FindNeigbhors and FindCluster function.

### Differential expression analysis and comparison of differentially expressed genes

Differential expression (DE) analysis was done between cells of interest with the FindMarkers function using the Wilcoxon Rank Sum test. Dissociation induced stress genes (van den Brin, SC, et al. Nat Meth. 2017) were removed from the list. For comparison between mouse genes and human genes, all mouse genes were converted to their human homologues using the R homologene package (1.4.68.19.3.27), for genes that did not have a human homologue, the case of the gene name was converted to capital. For comparisons between genes using different versions of the same gene name, gene lists were updated to their latest approved symbols by comparing to the Hugo Gene Nomenclature Committee (HGNC) database as of April 2023.

### Integrating datasets

To generate the integrated endothelial map, the pancreatic endothelial population from the Descartes fetal pancreas dataset, and the adult and neonatal pancreas dataset from Tosti at al. were merged. The endothelial cell subpopulation from the pancreas datasets were individually processed using the Seurat pipeline detailed above. After clustering, contaminating clusters were identified by comparing differentially expressed genes (DEGs) between clusters. Clusters with differentially upregulated non-vascular markers such as endocrine and exocrine pancreas cell markers were considered contaminating clusters. Once contaminating cells were removed, remaining endothelial cells from the different datasets were merged into one Seurat object using the merge function. The merged object was processed using the Seurat pipeline as detailed above, except the source of the dataset was regressed out while scaling the second time. The data was integrated with Harmony (v1.0) to remove variability from the source of the dataset (Descartes database or from Tosti. et. al.) followed by the individual donor. The resulting integrated or ‘harmonized’ principal components were used for UMAP dimension reduction and clustering as detailed above.

### Scorpius trajectory analysis

Pseudotime trajectory analysis was performed using the R package SCORPIUS. (v1.0.8). Trajectory inference was conducted on UMAP embeddings using the infer_trajectory function. For determining top genes involved in the ordering of the cells along pseudotime, top 50 markers were identified in each cluster in the integrated pancreas EC atlas through DE analysis as described above. Only the log-normalized counts matrix for these genes were retained to form a truncated expression matrix. SCORPIUS’s gene_importances function was then used to determine genes, whose expression level across the pseudotime best matches the time order using the gene_importances function.

### Bulk RNA-seq analysis

Log normalization of counts per million (CPM) was conducted on raw counts. Z-scores were obtained by scaling the LogCPM values using Seurat’s ScaleData function. Donor-specific batch affects were omitted in the scaled data by including ‘Donor’ as a variable to regress out during scaling. Principal component analysis was conducted using LogCPM values. Differential Expression analysis on exocrine and endocrine samples were conducted using DESeq2 using raw counts. Module score was derived using LogCPM values using the AddModuleScore function in Seurat.

### Data visualization

UMAPs, dot plots, box plots and volcano plots were generated using SCPubr (v1.1.2)(*91*). Hierarchical clustering were generated using ggtree(*92*) (v3.4.4) except for the heatmap for pseudotime trajectory analysis, which was generated using SCORPIUS’s draw_trajectory_heatmap function. Heatmap for bulk RNA-seq analysis was generated using heatmaply (1.4.2). Venn diagrams were plotted using ggvenn (0.1.9). Scatterplots were generated using ggplot2 (3.4.1). Breaks in violin plots were created using ggbreak (0.1.1)(*93*).

### Reference Mapping

To map a query dataset to the reference dataset, anchors were identified using Seurat’s FindTransferAnchors function and then the annotations from the reference dataset was projected onto the query using MapQuery

### *In situ* immunofluorescence staining

Tissue cross-sections were obtained from E18.5 mouse pancreatic tissue. Pregnant CD1 dams were euthanized via CO2 asphyxiation, in accordance with IACUC procedures at UT Southwestern Medical Center (UTSW). Embryos were dissected in ice cold -PBS, and pancreata were obtained for fresh-frozen embedding. Pancreata were briefly dabbed on a kimwipe, and washed 1x in Optimal Cutting Temperature (OCT) media. Pancreata were then placed in OCT-filled embedding and held over liquid nitrogen until frozen. Samples were then stored at −80C. Samples were cryosectioned by UTSW’s histology core. Briefly, slides were placed in 4% PFA at room temperature for an hour, and then wash 3X in PBS for 5 minutes per wash. Slides were permeabilized with 0.3% Triton for 12 minutes, and then placed in Cas Block (ThermoFisher Cat# 008120). The following primary antibodies were used: Insulin (Cell signaling technology Cat# 4590), Glucagon (Millipore Cat# 4030-01F), VE-Cadherin (Santa Cruz Cat# sc-6458), Pecam1 (Santa Cruz Cat# c-1506), Plvap (BD Pharmingen Cat# 550563). All antibodies were used at a 1:100 dilution, and slides were incubated in primary antibody overnight at 4C. Secondary antibodies conjugated to AlexaFlour were used for visualization.

### *In situ* RNAscope™ HiPlex v2 assay

Tissue cross-sections were obtained from E18.5 mouse pancreatic tissue. Pregnant CD1 dams were euthanized via CO2 asphyxiation, in accordance with IACUC procedures at UT Southwestern Medical Center (UTSW). For RNAscope™ HiPlex v2 assay (Advanced Cell Diagnostics), tissue was processed according to manufacturer’s instructions. Briefly, slides were fixed in 4% PFA for an hour. Slides were then washed 2x in PBS and dehydrated to 100% Ethanol. Slides were permeabilized with 12 minutes of protease III treatment. Target probes were applied at 1x concentration and incubated at 40 degrees for 2 hours. Amplification, fluorophore conjugation, and cleavage was done in accordance with manufacturers protocol. After all probes were imaged, slides were then cleaved 1 more time, washed three times in PBS for 5 minutes per wash. 150 µL of Cas Block was placed in on each slide for 1 hour at room temp, and then primary antibodies were added at a dilution of 1:100. Slides were then incubated in primary antibody overnight at 4°C Secondary antibodies conjugated to AlexaFlour were used for visualization.

### *In situ* Visualization

Images were taken on a nikon a1r confocal, a laser scanning system using the 40x objective (resolution of 3.2 pixels/ 1 µm). For RNA scope images, tile-scan images coupled with zstacks were taken to obtain large images of the pancreas section. Max Intensity projections were created in ImageJ, and then images were registered using ACD Hiplex Registration software.

### *In situ* image quantification

Images were quantified in a single-blind manner. 10 representative images were taken from the endocrine and exocrine region (based on staining of Insulin/Glucagon) of the capillary vessels based on size. Afterwards these images were analyzed on Fiji by a blinded researcher. RNA Probe channels for *Pecam1* and genes of interest were thresholded using the ‘MaxEntropy’ option and adjusted accordingly to remove background. Then regions of interest (ROI) was outlined manually based on the Pecam1/Endomucin staining channel. Afterwards, the area of probe was quantified and divided by the area of the ROIs to obtain normalized ratios of RNA probe to area of vessel.

### Statistical analysis

Differential expression analysis was conducted using the Wilcoxon Rank Sum Test. Afterwards the p-values obtained were adjusted using the Bonferroni correction method. For quantification of immunofluorescence staining and RNA probe binding, sample variances were tested for equality using var.equal function in R. If variances were equal then a one-tailed two-sample t-test was conducted, if variances were unequal then a Welch’s t-test was conducted. To compare bulk RNA-seq samples one-tailed students t-test was used on CPM normalized data.

### Data and Code Availability

All data are available in the main text or the supplementary materials. The code to recreate figures and analysis pipelines can be found at https://github.com/Safwat08/Pancecc/

## Acknowledgments, Sources of Funding, & Disclosures

### Acknowledgements

STK and SSN conceptualized the study. SSN obtained funding for the study. STK curated, processed, analyzed and visualized all single cell RNA sequencing data. STK and SV analyzed and visualized all Bulk RNA sequencing data. NA and OC performed the sample collection, staining, imaging and acquisition of all *in situ* data. ST quantified all *in situ* data. SSN supervised the study. STK and SSN wrote the original manuscript. ST, SV, NA, OC were involved in reviewing and editing the manuscript.

### Supplemental Material

Tables S1–S8

Figure S1-S8

## References

1. S. Rafii, J. M. Butler, B. Sen Ding, Angiocrine functions of organ-specific endothelial cells. Nature. 529, 316 (2016).

2. B. V. Zlokovic, The Blood-Brain Barrier in Health and Chronic Neurodegenerative Disorders. Neuron. 57 (2008), pp. 178–201.

3. J. Poisson, S. Lemoinne, C. Boulanger, F. Durand, R. Moreau, D. Valla, P. E. Rautou, Liver sinusoidal endothelial cells: Physiology and role in liver diseases. J Hepatol. 66 (2017), pp. 212–227.

4. D. J. Steiner, A. Kim, K. Miller, M. Hara, Pancreatic islet plasticity: Interspecies comparison of islet architecture and composition. Islets. 2, 135 (2010).

5. J. R. Henderson, M. C. Moss, A morphometric study of the endocrine and exocrine capillaries of the pancreas. Q J Exp Physiol. 70, 347–356 (1985).

6. P. Rorsman, F. M. Ashcroft, Pancreatic β-Cell Electrical Activity and Insulin Secretion: Of Mice and Men. Physiol Rev. 98, 117 (2018).

7. L. R. Nyman, E. Ford, A. C. Powers, D. W. Piston, Glucose-dependent blood flow dynamics in murine pancreatic islets in vivo. Am J Physiol Endocrinol Metab. 298, E807 (2010).

8. H. Peiris, C. S. Bonder, P. T. H. Coates, D. J. Keating, C. F. Jessup, The β-cell/EC axis: How do islet cells talk to each other? Diabetes. 63 (2014), pp. 3–11.

9. Å. Johansson, J. Lau, M. Sandberg, L. A. H. Borg, P. U. Magnusson, P. O. Carlsson, Endothelial cell signalling supports pancreatic beta cell function in the rat. Diabetologia. 52, 2385–2394 (2009).

10. R. Olsson, P. O. Carlsson, The pancreatic islet endothelial cell: Emerging roles in islet function and disease. Int J Biochem Cell Biol. 38, 710–714 (2006).

11. M. F. Hogan, D. J. Hackney, A. C. Aplin, T. O. Mundinger, M. J. Larmore, J. J. Castillo, N. Esser, S. Zraika, R. L. Hull, SGLT2-i improves markers of islet endothelial cell function in db/db diabetic mice. J Endocrinol. 248, 95–106 (2021).

12. J. C. Schupp, T. S. Adams, C. Cosme, M. S. B. Raredon, Y. Yuan, N. Omote, S. Poli, M. Chioccioli, K. A. Rose, E. P. Manning, M. Sauler, G. Deiuliis, F. Ahangari, N. Neumark, A. C. Habermann, A. J. Gutierrez, L. T. Bui, R. Lafyatis, R. W. Pierce, K. B. Meyer, M. C. Nawijn, S. A. Teichmann, N. E. Banovich, J. A. Kropski, L. E. Niklason, D. Pe’er, X. Yan, R. J. Homer, I. O. Rosas, N. Kaminski, Integrated Single-Cell Atlas of Endothelial Cells of the Human Lung. Circulation (2021), doi:10.1161/CIRCULATIONAHA.120.052318.

13. A. C. Yang, R. T. Vest, F. Kern, D. P. Lee, M. Agam, C. A. Maat, P. M. Losada, M. B. Chen, N. Schaum, N. Khoury, A. Toland, K. Calcuttawala, H. Shin, R. Pálovics, A. Shin, E. Y. Wang, J. Luo, D. Gate, W. J. Schulz-Schaeffer, P. Chu, J. A. Siegenthaler, M. W. McNerney, A. Keller, T. Wyss-Coray, A human brain vascular atlas reveals diverse mediators of Alzheimer’s risk. Nature. 603, 885 (2022).

14. M. Enge, H. E. Arda, M. Mignardi, J. Beausang, R. Bottino, S. K. Kim, S. R. Quake, Single-Cell Analysis of Human Pancreas Reveals Transcriptional Signatures of Aging and Somatic Mutation Patterns. Cell. 171, 321 (2017).

15. J. Camunas-Soler, X. Q. Dai, Y. Hang, A. Bautista, J. Lyon, K. Suzuki, S. K. Kim, S. R. Quake, P. E. MacDonald, Patch-seq links single-cell transcriptomes to human islet dysfunction in diabetes. Cell Metab. 31, 1017 (2020).

16. N. Lawlor, J. George, M. Bolisetty, R. Kursawe, L. Sun, V. Sivakamasundari, I. Kycia, P. Robson, M. L. Stitzel, Single-cell transcriptomes identify human islet cell signatures and reveal cell-type-specific expression changes in type 2 diabetes. Genome Res. 27, 208–222 (2017).

17. M. Baron, A. Veres, S. L. Wolock, A. L. Faust, R. Gaujoux, A. Vetere, J. H. Ryu, B. K. Wagner, S. S. Shen-Orr, A. M. Klein, D. A. Melton, I. Yanai, A Single-Cell Transcriptomic Map of the Human and Mouse Pancreas Reveals Inter- and Intra-cell Population Structure. Cell Syst. 3, 346–360.e4 (2016).

18. M. J. Muraro, G. Dharmadhikari, D. Grün, N. Groen, T. Dielen, E. Jansen, L. van Gurp, M. A. Engelse, F. Carlotti, E. J. P. de Koning, A. van Oudenaarden, A Single-Cell Transcriptome Atlas of the Human Pancreas. Cell Syst. 3, 385 (2016).

19. Å. Segerstolpe, A. Palasantza, P. Eliasson, E. M. Andersson, A. C. Andréasson, X. Sun, S. Picelli, A. Sabirsh, M. Clausen, M. K. Bjursell, D. M. Smith, M. Kasper, C. Ämmälä, R. Sandberg, Single-Cell Transcriptome Profiling of Human Pancreatic Islets in Health and Type 2 Diabetes. Cell Metab. 24, 593–607 (2016).

20. C. Ahlmann-Eltze, W. Huber, Comparison of transformations for single-cell RNA-seq data. Nature Methods 2023 20:5. 20, 665–672 (2023).

21. D. Wang, J. Wang, L. Bai, H. Pan, H. Feng, H. Clevers, Y. A. Zeng, Long-Term Expansion of Pancreatic Islet Organoids from Resident Procr+ Progenitors. Cell. 180, 1198–1211.e19 (2020).

22. R. B. Kang, Y. Li, C. Rosselot, T. Zhang, M. Siddiq, P. Rajbhandari, A. F. Stewart, D. K. Scott, A. Garcia-Ocana, G. Lu, Single-nucleus RNA sequencing of human pancreatic islets identifies novel gene sets and distinguishes β-cell subpopulations with dynamic transcriptome profiles. Genome Med. 15, 1–24 (2023).

23. J. Y. Chung, Y. Ma, D. Zhang, H. H. Bickerton, E. Stokes, S. B. Patel, H. M. Tse, J. Feduska, R. S. Welner, R. R. Banerjee, Pancreatic islet cell type–specific transcriptomic changes during pregnancy and postpartum. iScience. 26 (2023), doi:10.1016/j.isci.2023.106439.

24. B. Vieth, S. Parekh, C. Ziegenhain, W. Enard, I. Hellmann, A systematic evaluation of single cell RNA-seq analysis pipelines. Nat Commun. 10 (2019), doi:10.1038/S41467-019-12266-7.

25. J. Kalucka, L. P. M. H. de Rooij, J. Goveia, K. Rohlenova, S. J. Dumas, E. Meta, N. V. Conchinha, F. Taverna, L. A. Teuwen, K. Veys, M. García-Caballero, S. Khan, V. Geldhof, L. Sokol, R. Chen, L. Treps, M. Borri, P. de Zeeuw, C. Dubois, T. K. Karakach, K. D. Falkenberg, M. Parys, X. Yin, S. Vinckier, Y. Du, R. A. Fenton, L. Schoonjans, M. Dewerchin, G. Eelen, B. Thienpont, L. Lin, L. Bolund, X. Li, Y. Luo, P. Carmeliet, Single-Cell Transcriptome Atlas of Murine Endothelial Cells. Cell. 180, 764–779.e20 (2020).

26. D. T. Paik, L. Tian, I. M. Williams, S. Rhee, H. Zhang, C. Liu, R. Mishra, S. M. Wu, K. Red-Horse, J. C. Wu, Single-Cell RNA-seq Unveils Unique Transcriptomic Signatures of Organ-Specific Endothelial Cells. Circulation (2020), doi:10.1161/circulationaha.119.041433.

27. W. Feng, L. Chen, P. K. Nguyen, S. M. Wu, G. Li, Single Cell Analysis of Endothelial Cells Identified Organ-Specific Molecular Signatures and Heart-Specific Cell Populations and Molecular Features. Front Cardiovasc Med. 6, 165 (2019).

28. N. Schaum, J. Karkanias, N. F. Neff, A. P. May, S. R. Quake, T. Wyss-Coray, S. Darmanis, J. Batson, O. Botvinnik, M. B. Chen, S. Chen, F. Green, R. C. Jones, A. Maynard, L. Penland, A. O. Pisco, R. V. Sit, G. M. Stanley, J. T. Webber, F. Zanini, A. S. Baghel, I. Bakerman, I. Bansal, D. Berdnik, B. Bilen, D. Brownfield, C. Cain, M. Cho, G. Cirolia, S. D. Conley, A. Demers, K. Demir, A. de Morree, T. Divita, H. du Bois, L. B. T. Dulgeroff, H. Ebadi, F. H. Espinoza, M. Fish, Q. Gan, B. M. George, A. Gillich, G. Genetiano, X. Gu, G. S. Gulati, Y. Hang, S. Hosseinzadeh, A. Huang, T. Iram, T. Isobe, F. Ives, K. S. Kao, G. Karnam, A. M. Kershner, B. M. Kiss, W. Kong, M. E. Kumar, J. Y. Lam, D. P. Lee, S. E. Lee, G. Li, Q. Li, L. Liu, A. Lo, W. J. Lu, A. Manjunath, K. L. May, O. L. May, M. McKay, R. J. Metzger, M. Mignardi, D. Min, A. N. Nabhan, K. M. Ng, J. Noh, R. Patkar, W. C. Peng, R. Puccinelli, E. J. Rulifson, S. S. Sikandar, R. Sinha, K. Szade, W. Tan, C. Tato, K. Tellez, K. J. Travaglini, C. Tropini, L. Waldburger, L. J. van Weele, M. N. Wosczyna, J. Xiang, S. Xue, J. Youngyunpipatkul, M. E. Zardeneta, F. Zhang, L. Zhou, P. Castro, D. Croote, J. L. DeRisi, C. S. Kuo, B. Lehallier, P. K. Nguyen, S. Y. Tan, B. M. Wang, H. Yousef, P. A. Beachy, C. K. F. Chan, K. C. Huang, K. Weinberg, S. M. Wu, B. A. Barres, M. F. Clarke, S. K. Kim, M. A. Krasnow, R. Nusse, T. A. Rando, J. Sonnenburg, I. L. Weissman, Single-cell transcriptomics of 20 mouse organs creates a Tabula Muris. Nature. 562, 367–372 (2018).

29. J. Cao, D. R. O’Day, H. A. Pliner, P. D. Kingsley, M. Deng, R. M. Daza, M. A. Zager, K. A. Aldinger, R. Blecher-Gonen, F. Zhang, M. Spielmann, J. Palis, D. Doherty, F. J. Steemers, I. A. Glass, C. Trapnell, J. Shendure, A human cell atlas of fetal gene expression. Science (1979). 370 (2020), doi:10.1126/SCIENCE.ABA7721/SUPPL_FILE/ABA7721_TABLESS1-S16.XLSX.

30. R. C. Jones, J. Karkanias, M. A. Krasnow, A. O. Pisco, S. R. Quake, J. Salzman, N. Yosef, B. Bulthaup, P. Brown, W. Harper, M. Hemenez, R. Ponnusamy, A. Salehi, B. A. Sanagavarapu, E. Spallino, K. A. Aaron, W. Concepcion, J. M. Gardner, B. Kelly, N. Neidlinger, Z. Wang, S. Crasta, S. Kolluru, M. Morri, A. O. Pisco, S. Y. Tan, K. J. Travaglini, C. Xu, M. Alcántara-Hernández, N. Almanzar, J. Antony, B. Beyersdorf, D. Burhan, K. Calcuttawala, M. M. Carter, C. K. F. Chan, C. A. Chang, S. Chang, A. Colville, S. Crasta, R. N. Culver, I. Cvijović, G. D’Amato, C. Ezran, F. X. Galdos, A. Gillich, W. R. Goodyer, Y. Hang, A. Hayashi, S. Houshdaran, X. Huang, J. C. Irwin, S. Jang, J. V. Juanico, A. M. Kershner, S. Kim, B. Kiss, S. Kolluru, W. Kong, M. E. Kumar, A. H. Kuo, R. Leylek, B. Li, G. B. Loeb, W.-J. Lu, S. Mantri, M. Markovic, P. L. McAlpine, A. de Morree, M. Morri, K. Mrouj, S. Mukherjee, T. Muser, P. Neuhöfer, T. D. Nguyen, K. Perez, R. Phansalkar, A. O. Pisco, N. Puluca, Z. Qi, P. Rao, H. Raquer-McKay, N. Schaum, B. Scott, B. Seddighzadeh, J. Segal, S. Sen, S. Sikandar, S. P. Spencer, L. C. Steffes, V. R. Subramaniam, A. Swarup, M. Swift, K. J. Travaglini, W. Van Treuren, E. Trimm, S. Veizades, S. Vijayakumar, K. C. Vo, S. K. Vorperian, W. Wang, H. N. W. Weinstein, J. Winkler, T. T. H. Wu, J. Xie, A. R. Yung, Y. Zhang, A. M. Detweiler, H. Mekonen, N. F. Neff, R. V. Sit, M. Tan, J. Yan, G. R. Bean, V. Charu, E. Forgó, B. A. Martin, M. G. Ozawa, O. Silva, S. Y. Tan, A. Toland, V. N. P. Vemuri, S. Afik, K. Awayan, O. B. Botvinnik, A. Byrne, M. Chen, R. Dehghannasiri, A. M. Detweiler, A. Gayoso, A. A. Granados, Q. Li, G. Mahmoudabadi, A. McGeever, A. de Morree, J. E. Olivieri, M. Park, A. O. Pisco, N. Ravikumar, J. Salzman, G. Stanley, M. Swift, M. Tan, W. Tan, A. J. Tarashansky, R. Vanheusden, S. K. Vorperian, P. Wang, S. Wang, G. Xing, C. Xu, N. Yosef, M. Alcántara-Hernández, J. Antony, C. K. F. Chan, C. A. Chang, A. Colville, S. Crasta, R. Culver, L. Dethlefsen, C. Ezran, A. Gillich, Y. Hang, P.-Y. Ho, J. C. Irwin, S. Jang, A. M. Kershner, W. Kong, M. E. Kumar, A. H. Kuo, R. Leylek, S. Liu, G. B. Loeb, W.-J. Lu, J. S. Maltzman, R. J. Metzger, A. de Morree, P. Neuhöfer, K. Perez, R. Phansalkar, Z. Qi, P. Rao, H. Raquer-McKay, K. Sasagawa, B. Scott, R. Sinha, H. Song, S. P. Spencer, A. Swarup, M. Swift, K. J. Travaglini, E. Trimm, S. Veizades, S. Vijayakumar, B. Wang, W. Wang, J. Winkler, J. Xie, A. R. Yung, S. E. Artandi, P. A. Beachy, M. F. Clarke, L. C. Giudice, F. W. Huang, K. C. Huang, J. Idoyaga, S. K. Kim, M. Krasnow, C. S. Kuo, P. Nguyen, S. R. Quake, T. A. Rando, K. Red-Horse, J. Reiter, D. A. Relman, J. L. Sonnenburg, B. Wang, A. Wu, S. M. Wu, T. Wyss-Coray, The Tabula Sapiens: A multiple-organ, single-cell transcriptomic atlas of humans. Science (1979). 376 (2022), doi:10.1126/SCIENCE.ABL4896/SUPPL_FILE/SCIENCE.ABL4896_MDAR_REPRODUCIBILITY_CHECKLIST.PDF.

31. B. Marquina-Sanchez, N. Fortelny, M. Farlik, A. Vieira, P. Collombat, C. Bock, S. Kubicek, Single-cell RNA-seq with spike-in cells enables accurate quantification of cell-specific drug effects in pancreatic islets. Genome Biol. 21 (2020), doi:10.1186/S13059-020-02006-2.

32. M. D. Young, S. Behjati, SoupX removes ambient RNA contamination from droplet-based single-cell RNA sequencing data. Gigascience. 9, 1–10 (2020).

33. S. Yang, S. E. Corbett, Y. Koga, Z. Wang, W. E. Johnson, M. Yajima, J. D. Campbell, Decontamination of ambient RNA in single-cell RNA-seq with DecontX. Genome Biol. 21 (2020), doi:10.1186/S13059-020-1950-6.

34. L. Tosti, Y. Hang, O. Debnath, S. Tiesmeyer, T. Trefzer, K. Steiger, F. W. Ten, S. Lukassen, S. Ballke, A. A. Kühl, S. Spieckermann, R. Bottino, N. Ishaque, W. Weichert, S. K. Kim, R. Eils, C. Conrad, Single-Nucleus and In Situ RNA–Sequencing Reveal Cell Topographies in the Human Pancreas. Gastroenterology. 160, 1330–1344.e11 (2021).

35. Y. Hao, S. Hao, E. Andersen-Nissen, W. M. Mauck, S. Zheng, A. Butler, M. J. Lee, A. J. Wilk, C. Darby, M. Zager, P. Hoffman, M. Stoeckius, E. Papalexi, E. P. Mimitou, J. Jain, A. Srivastava, T. Stuart, L. M. Fleming, B. Yeung, A. J. Rogers, J. M. McElrath, C. A. Blish, R. Gottardo, P. Smibert, R. Satija, Integrated analysis of multimodal single-cell data. Cell. 184, 3573 (2021).

36. I. Tirosh, B. Izar, S. M. Prakadan, M. H. Wadsworth, D. Treacy, J. J. Trombetta, A. Rotem, C. Rodman, C. Lian, G. Murphy, M. Fallahi-Sichani, K. Dutton-Regester, J. R. Lin, O. Cohen, P. Shah, D. Lu, A. S. Genshaft, T. K. Hughes, C. G. K. Ziegler, S. W. Kazer, A. Gaillard, K. E. Kolb, A. C. Villani, C. M. Johannessen, A. Y. Andreev, E. M. Van Allen, M. Bertagnolli, P. K. Sorger, R. J. Sullivan, K. T. Flaherty, D. T. Frederick, J. Jané-Valbuena, C. H. Yoon, O. Rozenblatt-Rosen, A. K. Shalek, A. Regev, L. A. Garraway, Dissecting the multicellular ecosystem of metastatic melanoma by single-cell RNA-seq. Science. 352, 189 (2016).

37. H. C. Wang, T. H. Wong, L. T. Wang, H. H. Su, H. Y. Yu, A. H. Wu, Y. C. Lin, H. L. Chen, J. L. Suen, S. H. Hsu, L. C. Chen, Y. Zhou, S. K. Huang, Aryl hydrocarbon receptor signaling promotes ORMDL3-dependent generation of sphingosine-1-phosphate by inhibiting sphingosine-1-phosphate lyase. Cell Mol Immunol. 16, 783 (2019).

38. N. R. Reinhard, M. Mastop, T. Yin, Y. Wu, E. K. Bosma, T. W. J. Gadella, J. Goedhart, P. L. Hordijk, The balance between Gαi-Cdc42/Rac and Gα12/13-RhoA pathways determines endothelial barrier regulation by sphingosine-1-phosphate. Mol Biol Cell. 28, 3371 (2017).

39. J. G. N. Garcia, F. Liu, A. D. Verin, A. Birukova, M. A. Dechert, W. T. Gerthoffer, J. R. Bamburg, D. English, Sphingosine 1-phosphate promotes endothelial cell barrier integrity by Edg-dependent cytoskeletal rearrangement. J Clin Invest. 108, 689–701 (2001).

40. H. Chen, S. Ahmed, H. Zhao, N. Elghobashi-Meinhardt, Y. Dai, J. H. Kim, J. G. McDonald, X. Li, C. H. Lee, Structural and functional insights into Spns2-mediated transport of sphingosine-1-phosphate. Cell. 186, 2644–2655.e16 (2023).

41. H. Goto, M. Miyamoto, A. Kihara, Direct uptake of sphingosine-1-phosphate independent of phospholipid phosphatases. Journal of Biological Chemistry. 296, 100605 (2021).

42. I. Korsunsky, N. Millard, J. Fan, K. Slowikowski, F. Zhang, K. Wei, Y. Baglaenko, M. Brenner, P. ru Loh, S. Raychaudhuri, Fast, sensitive and accurate integration of single-cell data with Harmony. Nat Methods. 16, 1289–1296 (2019).

43. H. C. T. Nguyen, B. Baik, S. Yoon, T. Park, D. Nam, Benchmarking integration of single-cell differential expression. Nature Communications 2023 14:1. 14, 1–16 (2023).

44. I. Buschmann, A. Pries, B. Styp-Rekowska, P. Hillmeister, L. Loufrani, D. Henrion, Y. Shi, A. Duelsner, I. Hoefer, N. Gatzke, H. Wang, K. Lehmann, L. Ulm, Z. Ritter, P. Hauff, R. Hlushchuk, V. Djonov, T. Van Veen, F. Le Noble, Pulsatile shear and Gja5 modulate arterial identity and remodeling events during flow-driven arteriogenesis. Development. 137, 2187–2196 (2010).

45. J. S. Fang, B. G. Coon, N. Gillis, Z. Chen, J. Qiu, T. W. Chittenden, J. M. Burt, M. A. Schwartz, K. K. Hirschi, Shear-induced Notch-Cx37-p27 axis arrests endothelial cell cycle to enable arterial specification. Nature Communications 2017 8:1. 8, 1–14 (2017).

46. D. Shin, G. Garcia-Cardena, S. I. Hayashi, S. Gerety, T. Asahara, G. Stavrakis, J. Isner, J. Folkman, M. A. Gimbrone, D. J. Anderson, Expression of ephrinB2 identifies a stable genetic difference between arterial and venous vascular smooth muscle as well as endothelial cells, and marks subsets of microvessels at sites of adult neovascularization. Dev Biol. 230, 139–150 (2001).

47. B. Gorsi, F. Liu, X. Ma, T. J. A. Chico, A. Shrinivasan, K. L. Kramer, E. Bridges, R. Monteiro, A. L. Harris, R. Patient, S. E. Stringer, The heparan sulfate editing enzyme Sulf1 plays a novel role in zebrafish VegfA mediated arterial venous identity. Angiogenesis. 17, 77–91 (2014).

48. A. Thiriot, C. Perdomo, G. Cheng, I. Novitzky-Basso, S. McArdle, J. K. Kishimoto, O. Barreiro, I. Mazo, R. Triboulet, K. Ley, A. Rot, U. H. von Andrian, Differential DARC/ACKR1 expression distinguishes venular from non-venular endothelial cells in murine tissues. BMC Biol. 15, 1–19 (2017).

49. R. V. Stan, M. Kubitza, G. E. Palade, PV-1 is a component of the fenestral and stomatal diaphragms in fenestrated endothelia. Proc Natl Acad Sci U S A. 96, 13203 (1999).

50. C. G. Hansen, N. A. Bright, G. Howard, B. J. Nichols, SDPR induces membrane curvature and functions in the formation of caveolae. Nature Cell Biology 2009 11:7. 11, 807–814 (2009).

51. I. Yeow, G. Howard, J. Chadwick, C. Mendoza-Topaz, C. G. Hansen, B. J. Nichols, E. Shvets, EHD Proteins Cooperate to Generate Caveolar Clusters and to Maintain Caveolae during Repeated Mechanical Stress. Current Biology. 27, 2951 (2017).

52. N. Gil-Yarom, L. Radomir, L. Sever, M. P. Kramer, H. Lewinsky, C. Bornstein, R. Blecher-Gonen, Z. Barnett-Itzhaki, V. Mirkin, G. Friedlander, L. Shvidel, Y. Herishanu, E. J. Lolis, S. Becker-Herman, I. Amit, I. Shachar, CD74 is a novel transcription regulator. Proc Natl Acad Sci U S A. 114, 562 (2017).

53. S. M. Moore, V. V. Holt, L. R. Malpass, I. N. Hines, M. D. Wheeler, Fatty acid-binding protein 5 limits the anti-inflammatory response in murine macrophages. Mol Immunol. 67, 265 (2015).

54. R. Alejandro, F. L. Shienvold, S. Vaerewyck Hajek, U. Ryan, J. Miller, D. H. Mintz, Immunocytochemical Localization of HLA-DR in Human Islets of Langerhans. Diabetes. 31, 17–22 (1982).

55. W. Saelens, R. Cannoodt, H. Todorov, Y. Saeys, A comparison of single-cell trajectory inference methods. Nat Biotechnol. 37, 547–554 (2019).

56. K. Boyé, L. H. Geraldo, J. Furtado, L. Pibouin-Fragner, M. Poulet, D. Kim, B. Nelson, Y. Xu, L. Jacob, N. Maissa, D. Agalliu, L. Claesson-Welsh, S. L. Ackerman, A. Eichmann, Endothelial Unc5B controls blood-brain barrier integrity. Nature Communications 2022 13:1. 13, 1–15 (2022).

57. A. Jonsson, A. Hedin, M. Müller, O. Skog, O. Korsgren, Transcriptional profiles of human islet and exocrine endothelial cells in subjects with or without impaired glucose metabolism. Sci Rep. 10, 22315 (2020).

58. Y. Aghazadeh, S. T. Khan, B. Nkennor, S. S. Nunes, Cell-based therapies for vascular regeneration: Past, present and future. Pharmacol Ther. 231, 107976 (2022).

59. Y. Aghazadeh, F. Poon, F. Sarangi, F. T. M. Wong, S. T. Khan, X. Sun, R. Hatkar, B. J. Cox, S. S. Nunes, M. C. Nostro, Microvessels support engraftment and functionality of human islets and hESC-derived pancreatic progenitors in diabetes models. Cell Stem Cell. 28, 1936–1949.e8 (2021).

60. S. S. Hasan, M. Jabs, J. Taylor, L. Wiedmann, T. Leibing, V. Nordström, G. Federico, L. P. Roma, C. Carlein, G. Wolff, B. Ekim-Üstünel, M. Brune, I. Moll, F. Tetzlaff, H.-J. Gröne, T. Fleming, C. Géraud, S. Herzig, P. P. Nawroth, A. Fischer, Endothelial Notch signaling controls insulin transport in muscle. EMBO Mol Med. 12, e09271 (2020).

61. J. Gustavsson, S. Parpal, P. Strålfors, Insulin-stimulated glucose uptake involves the transition of glucose transporters to a caveolae-rich fraction within the plasma membrane: Implications for type II diabetes. Molecular Medicine. 2, 367–372 (1996).

62. H. Wang, A. X. Wang, E. J. Barrett, Caveolin-1 is required for vascular endothelial insulin uptake. Am J Physiol Endocrinol Metab. 300, E134 (2011).

63. S. Hayashi, M. P. Scott, What determines the specificity of action of Drosophila homeodomain proteins? Cell. 63, 883–894 (1990).

64. M. N. Stanfel, K. A. Moses, R. J. Schwartz, W. E. Zimmer, Regulation of organ development by the NKX-homeodomain factors: an NKX code. Cell Mol Biol (Noisy-le-grand). Suppl 51, OL785-99 (2005).

65. O. Pabst, R. Zweigerdt, H. H. Arnold, Targeted disruption of the homeobox transcription factor Nkx2-3 in mice results in postnatal lethality and abnormal development of small intestine and spleen. Development. 126, 2215–2225 (1999).

66. T. Czömpöly, Á. Lábadi, Z. Kellermayer, K. Olasz, H.-H. Arnold, P. Balogh, Transcription Factor Nkx2-3 Controls the Vascular Identity and Lymphocyte Homing in the Spleen. The Journal of Immunology. 186, 6981–6989 (2011).

67. T. T. Dinh, M. Xiang, A. Rajaraman, Y. Wang, N. Salazar, Y. Zhu, W. Roper, S. Rhee, K. Brulois, E. O’Hara, H. Kiefel, T. M. Dinh, Y. Bi, D. Gonzalez, E. P. Bao, K. Red-Horse, P. Balogh, F. Gábris, B. Gaszner, G. Berta, J. Pan, E. C. Butcher, An NKX-COUP-TFII morphogenetic code directs mucosal endothelial addressin expression. Nature Communications 2022 13:1. 13, 1–14 (2022).

68. M. Nakache, E. Lakey Berg, P. R. Streeter, E. C. Butcher, The mucosal vascular addressin is a tissue-specific endothelial cell adhesion molecule for circulating lymphocytes. Nature 1989 337:6203. 337, 179–181 (1989).

69. M. Briskin, D. Winsor-Hines, A. Shyjan, N. Cochran, S. Bloom, J. Wilson, L. M. McEvoy, E. C. Butcher, N. Kassam, C. R. Mackay, W. Newman, D. J. Ringler, Human mucosal addressin cell adhesion molecule-1 is preferentially expressed in intestinal tract and associated lymphoid tissue. Am J Pathol. 151, 97 (1997).

70. W. Yu, J. P. Hegarty, A. Berg, X. Chen, G. West, A. A. Kelly, Y. Wang, L. S. Poritz, W. A. Koltun, Z. Lin, NKX2-3 Transcriptional Regulation of Endothelin-1 and VEGF Signaling in Human Intestinal Microvascular Endothelial Cells. PLoS One. 6, e20454 (2011).

71. N. Ancellin, C. Colmont, J. Su, Q. Li, N. Mittereder, S.-S. Chae, S. Stefansson, G. Liau, T. Hla, Extracellular Export of Sphingosine Kinase-1 Enzyme. Journal of Biological Chemistry. 277, 6667–6675 (2002).

72. A. Cartier, T. Hla, Sphingosine 1-phosphate: Lipid signaling in pathology and therapy. Science (1979). 366 (2019), doi:10.1126/SCIENCE.AAR5551/ASSET/C37AF20A-7636-40C9-A49A-4FC785158A19/ASSETS/GRAPHIC/366_AAR5551_FA.JPEG.

73. J. Edsbagge, J. K. Johansson, F. Esni, Y. Luo, G. L. Radice, H. Semb, Vascular function and sphingosine-1-phosphate regulate development of the dorsal pancreatic mesenchyme. Development. 132, 1085–1092 (2005).

74. J. C. Stanford, A. J. Morris, M. Sunkara, G. J. Popa, K. L. Larson, S. Özcan, Sphingosine 1-Phosphate (S1P) Regulates Glucose-stimulated Insulin Secretion in Pancreatic Beta Cells. J Biol Chem. 287, 13457 (2012).

75. H. Shimizu, F. Okajima, T. Kimura, K. I. Ohtani, T. Tsuchiya, H. Takahashi, A. Kuwabara, H. Tomura, K. Sato, M. Mori, Sphingosine 1-phosphate stimulates insulin secretion in HIT-T 15 cells and mouse islets. Endocr J. 47, 261–269 (2000).

76. Y. He, B. Shi, X. Zhao, J. Sui, Sphingosine-1-phosphate induces islet β-cell proliferation and decreases cell apoptosis in high-fat diet/streptozotocin diabetic mice. Exp Ther Med. 18, 3415 (2019).

77. P. M. Fernandez-Salguero, J. M. Ward, J. P. Sundberg, F. J. Gonzalez, Lesions of Aryl-hydrocarbon Receptor–deficient Mice. 10.1177/030098589703400609. 34, 605–614 (1997).

78. S. Ichihara, Y. Yamada, G. Ichihara, T. Nakajima, P. Li, T. Kondo, F. J. Gonzalez, T. Murohara, A role for the aryl hydrocarbon receptor in regulation of ischemia-induced angiogenesis. Arterioscler Thromb Vasc Biol. 27, 1297–1304 (2007).

79. S. Majumder, M. Kono, Y. Terry Lee, C. Byrnes, C. Li, G. Tuymetova, R. L. Proia, A genome-wide CRISPR/Cas9 screen reveals that the aryl hydrocarbon receptor stimulates sphingolipid levels. J Biol Chem. 295, 4341–4349 (2020).

80. D. M. Adyshev, N. K. Moldobaeva, V. R. Elangovan, J. G. N. Garcia, S. M. Dudek, Differential involvement of ezrin/radixin/moesin proteins in sphingosine 1-phosphate-induced human pulmonary endothelial cell barrier enhancement. Cell Signal. 23, 2086 (2011).

81. R. A. Brekken, E. H. Sage, SPARC, a matricellular protein: at the crossroads of cell-matrix. Matrix Biol. 19, 569–580 (2000).

82. C. L. Ryall, K. Viloria, F. Lhaf, A. J. Walker, A. King, P. Jones, D. Mackintosh, R. McNeice, H. Kocher, M. Flodstrom-Tullberg, C. Edling, N. J. Hill, Novel role for matricellular proteins in the regulation of islet β cell survival: the effect of SPARC on survival, proliferation, and signaling. J Biol Chem. 289, 30614–30624 (2014).

83. L. Hu, F. He, M. Huang, Q. Zhao, L. Cheng, N. Said, Z. Zhou, F. Liu, Y. S. Dai, SPARC promotes insulin secretion through down-regulation of RGS4 protein in pancreatic β cells. Sci Rep. 10, 17581 (2020).

84. C. Atorrasagasti, A. Onorato, M. L. Gimeno, L. Andreone, M. Garcia, M. Malvicini, E. Fiore, J. Bayo, M. J. Perone, G. D. Mazzolini, SPARC is required for the maintenance of glucose homeostasis and insulin secretion in mice. Clin Sci (Lond). 133, 351–365 (2019).

85. L. W. Harries, L. J. McCulloch, J. E. Holley, T. J. Rawling, H. J. Welters, K. Kos, A Role for SPARC in the Moderation of Human Insulin Secretion. PLoS One. 8, e68253 (2013).

86. L. Xu, F. Ping, J. Yin, X. Xiao, H. Xiang, C. M. Ballantyne, H. Wu, M. Li, Elevated Plasma SPARC Levels Are Associated with Insulin Resistance, Dyslipidemia, and Inflammation in Gestational Diabetes Mellitus. PLoS One. 8, 81615 (2013).

87. H. O. Steinberg, G. Brechtel, A. Johnson, N. Fineberg, A. D. Baron, Insulin-mediated skeletal muscle vasodilation is nitric oxide dependent. A novel action of insulin to increase nitric oxide release. Journal of Clinical Investigation. 94, 1172 (1994).

88. J. Almaça, J. Weitz, R. Rodriguez-Diaz, E. Pereira, A. Caicedo, THE PERICYTE OF THE PANCREATIC ISLET REGULATES CAPILLARY DIAMETER AND LOCAL BLOOD FLOW. Cell Metab. 27, 630 (2018).

89. M. D. Luecken, M. Büttner, K. Chaichoompu, A. Danese, M. Interlandi, M. F. Mueller, D. C. Strobl, L. Zappia, M. Dugas, M. Colomé-Tatché, F. J. Theis, Benchmarking atlas-level data integration in single-cell genomics. Nature Methods 2021 19:1. 19, 41–50 (2021).

90. H. T. N. Tran, K. S. Ang, M. Chevrier, X. Zhang, N. Y. S. Lee, M. Goh, J. Chen, A benchmark of batch-effect correction methods for single-cell RNA sequencing data. Genome Biol. 21, 1–32 (2020).

91. E. Blanco-Carmona, bioRxiv, in press, doi:10.1101/2022.02.28.482303.

92. G. Yu, D. K. Smith, H. Zhu, Y. Guan, T. T. Y. Lam, ggtree: an r package for visualization and annotation of phylogenetic trees with their covariates and other associated data. Methods Ecol Evol. 8, 28–36 (2017).

93. S. Xu, M. Chen, T. Feng, L. Zhan, L. Zhou, G. Yu, Use ggbreak to Effectively Utilize Plotting Space to Deal With Large Datasets and Outliers. Front Genet. 12, 2122 (2021).

