## Supplemental figures for "Uncovering the heterogeneity of pancreatic endothelial cells using integrative and comparative single cell gene expression analysis"

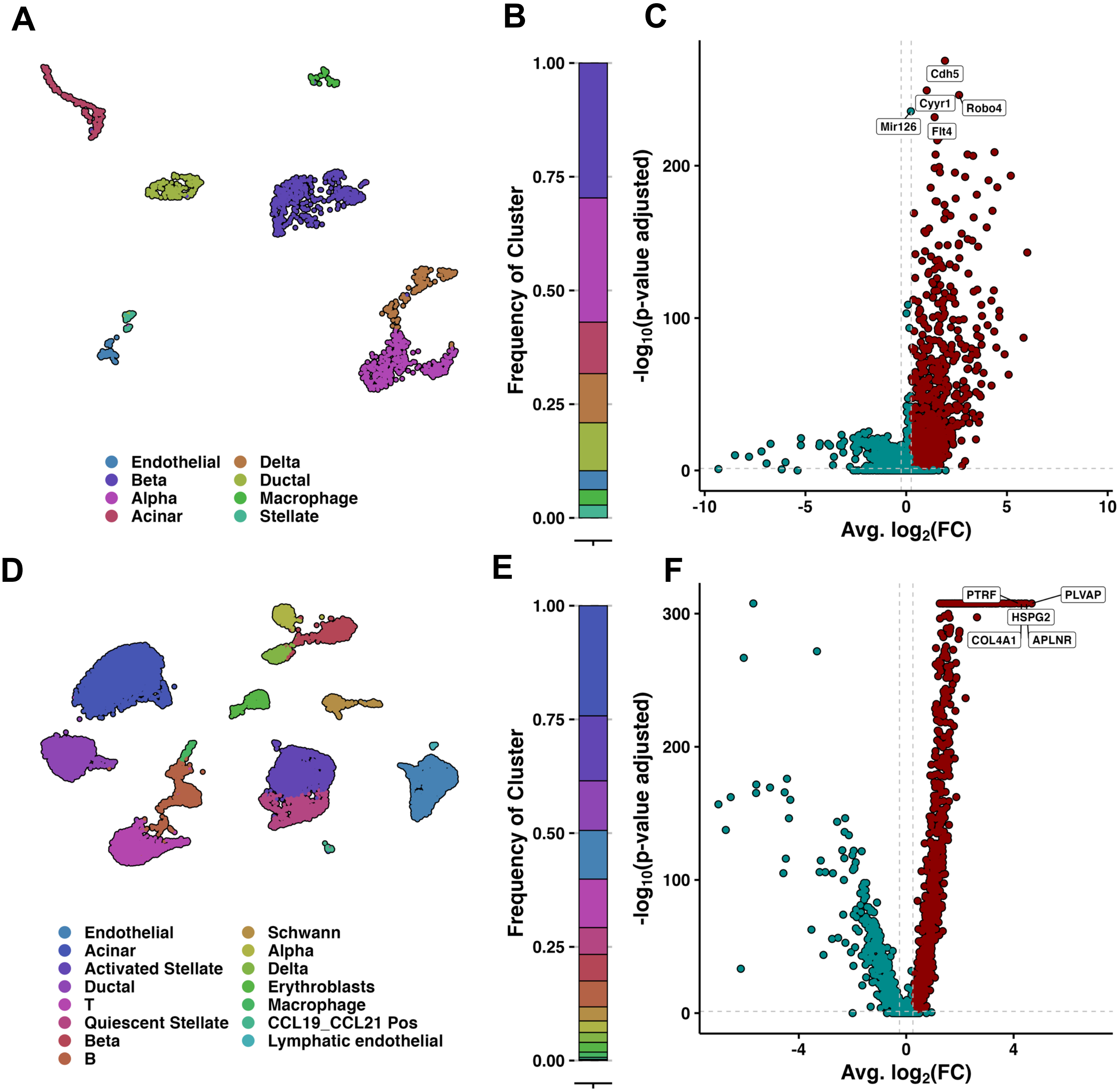


**Supplementary Figure 1. Summary of pancreas datasets derived from the mutli-organ gene expression atlases.** This shows the summary of pancreas datasets for adult mouse, the Tabula Muris (A-C) and fetal human, Descartes (D-F). A) and D) shows The UMAP visualization of single cell sequencing data. B and E) Highlight the the corresponding cell-type ratios in the pancreas datasets, the colors correspond to the labelling in the UMAPs. C and F) are volcano plot demonstrating differentially expressed genes (DEGs) in vascular ECs against other cell-types except lymphatic ECs and stellate cells. Significant DEGs (log2FoldChange > 0.25; Wilcoxon Rank Sum Test, Bonferroni-adjusted p-value < 0.05) are shown in red and the rest of the DEGs in blue. Further, the top 5 significant DEGs are labeled.

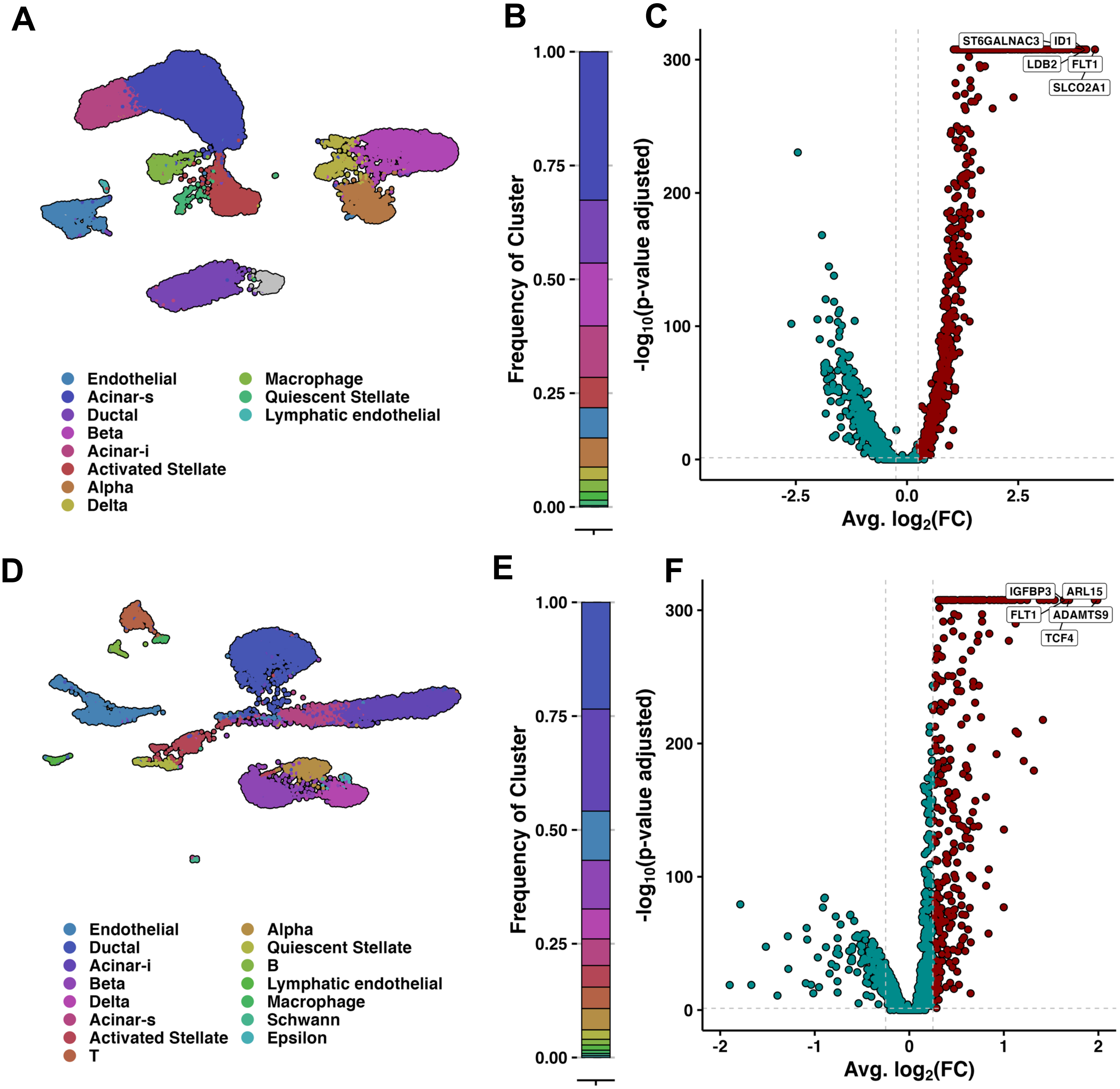


**Supplementary Figure 2. Summary of pancreas datasets from Tosti et al.** This shows the summary of pancreas datasets derived from Tosti et al. for adult human (A-C) and neonatl human (D-F). A) and D) shows The UMAP visualization of single cell sequencing data. B and E) highlight the the corresponding cell-type ratios in the pancreas datasets, the colors correspond to the labelling in the UMAPs. C and F) are volcano plot demonstrating differentially expressed genes (DEGs) in vascular ECs against other cell-types except lymphatic ECs and stellate cells. Significant DEGs (log2FoldChange > 0.25; Wilcoxon Rank Sum Test, Bonferroni-adjusted p-value < 0.05) are shown in red and the rest of the DEGs in blue. Further, the top 5 significant DEGs are labeled.

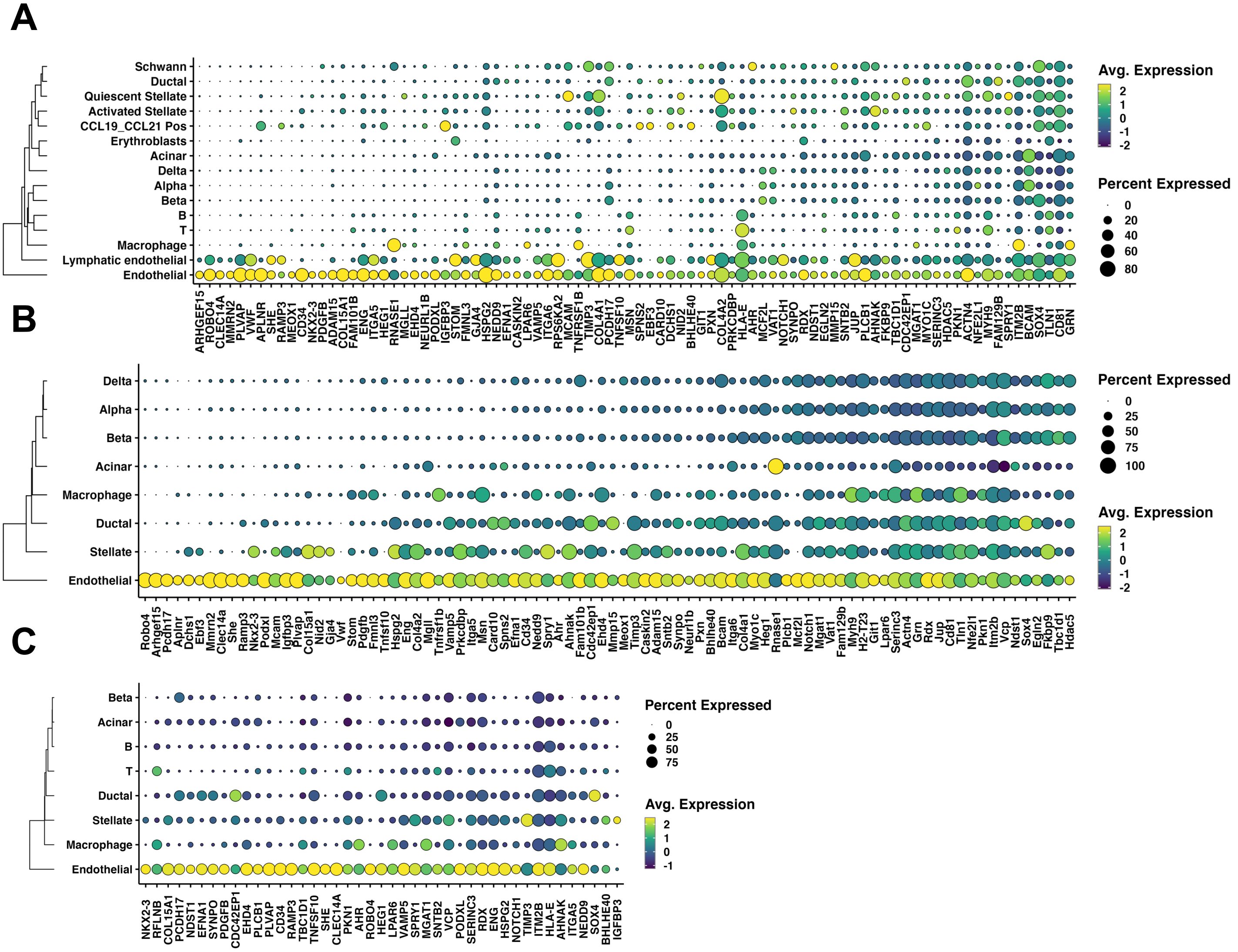


**Supplementary Figure 3. Assessment of the pancreas-specific EC gene signature across pancreas datasets obtained from all multi-organ gene expression atlases.** A-C) demonstrates the individual log-normalized average scaled expression of the pancreas-specific EC signature genes in the pancreas dataset from A) Descartes B) Tabula Muris and C) Tabula Sapiens atlas. Hierarchical clustering based on Euclidian distances between the average scaled expression of the DEGs across the different tissues demonstrates specificity to the pancreas.

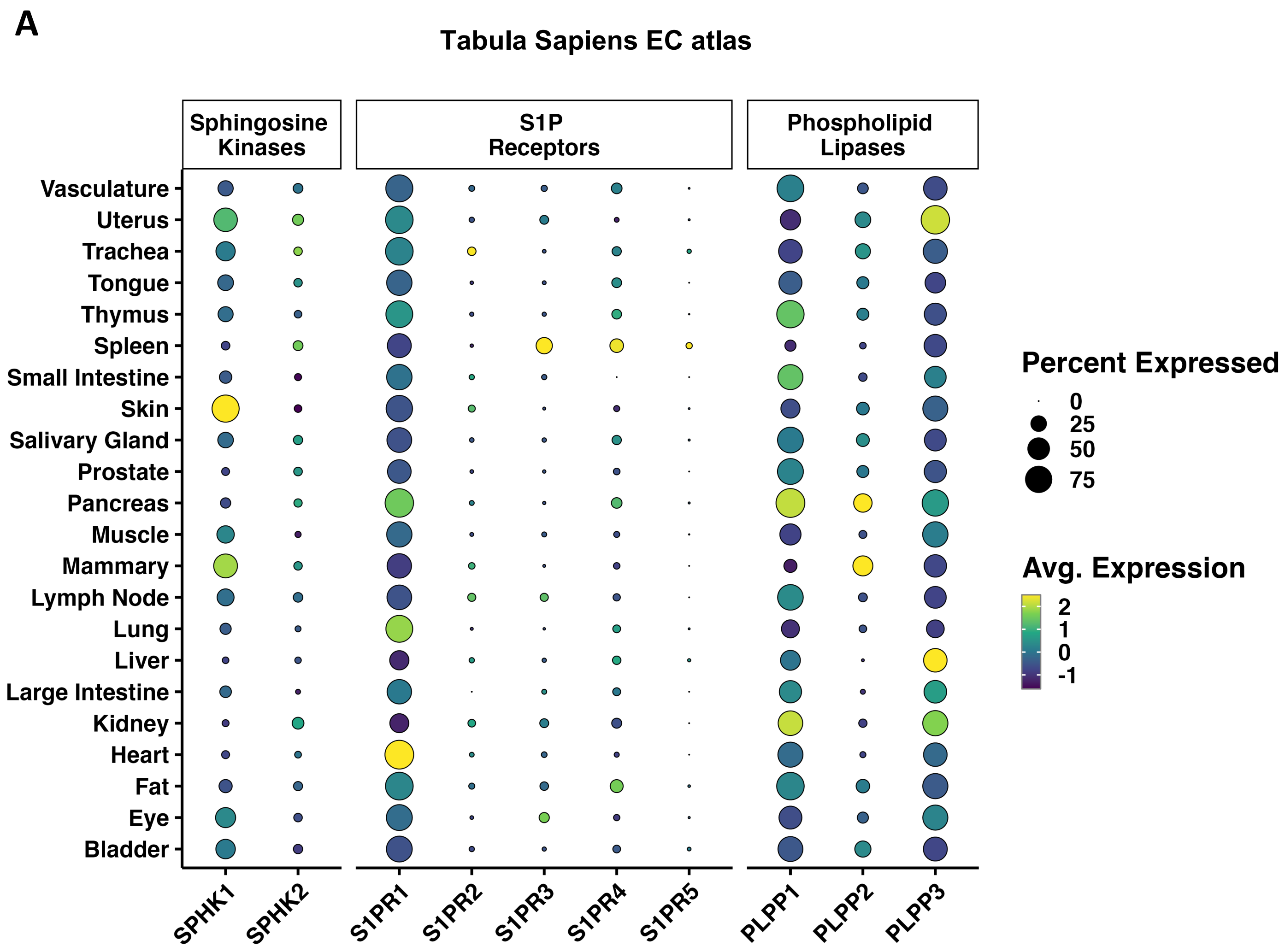


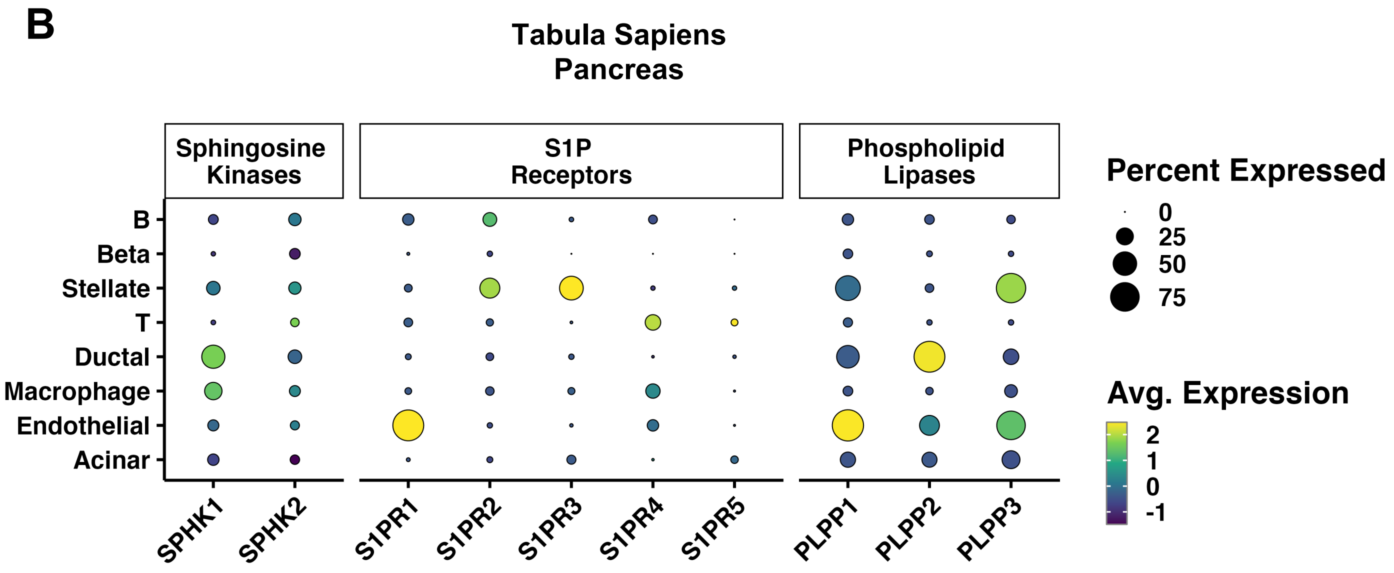


**Supplementary Figure 4. Assessment of sphingosine-1-phosphate signalling related genes in the Tabula Sapiens EC atlas.** Demonstrates the average expression of key sphingosine-1-phosphate associated molecules in A) the Tabula Sapiens EC atlas and in B) the Tabula Sapiens pancreas dataset.

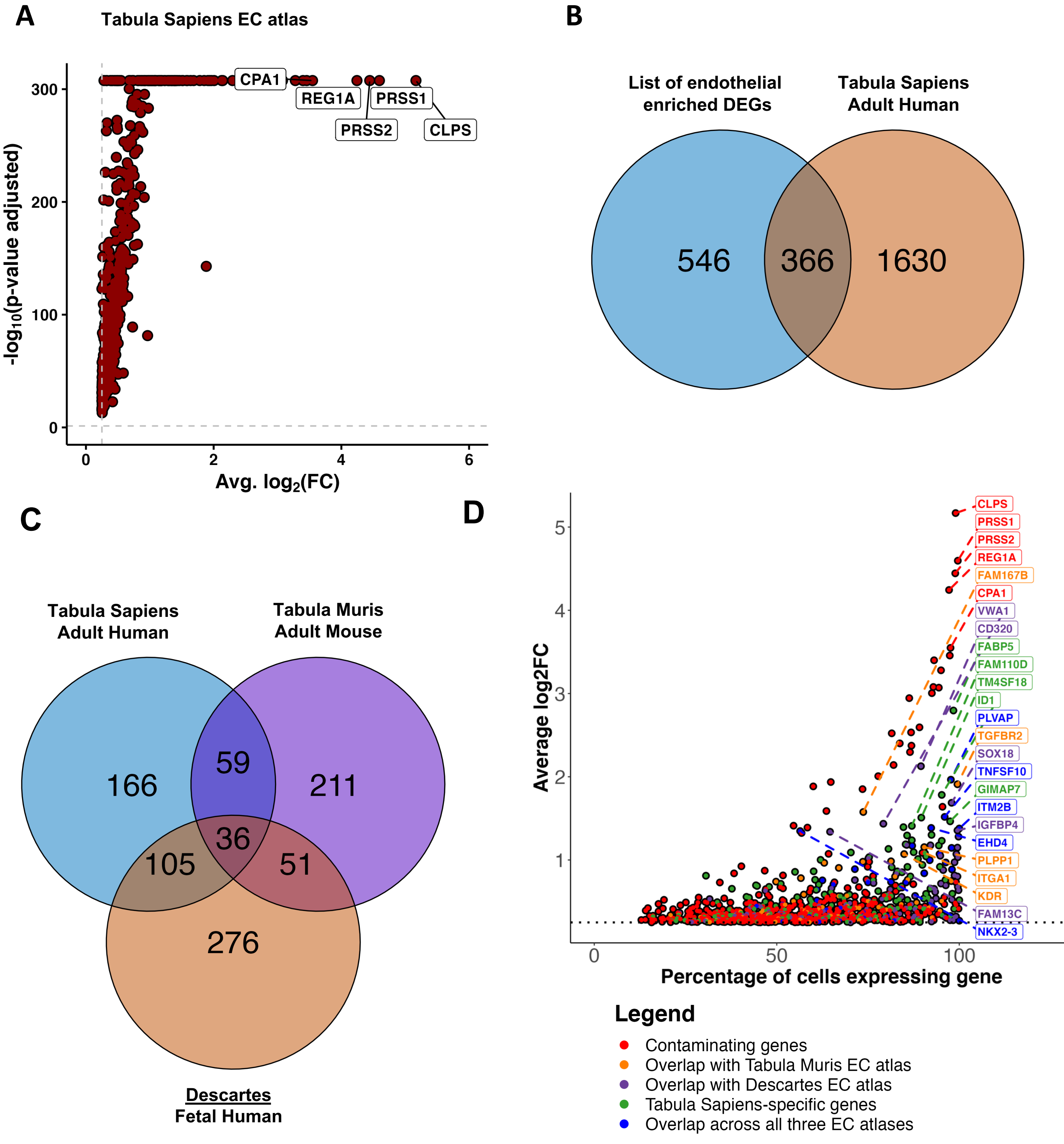


**Supplementary Figure 5. Assessment of the pancreas-enriched differentially expressed genes in the Tabula Sapiens endothelial cell (EC) atlas.** A) Demonstrates volcano plots for the Tabula Sapiens atlas, highlighting only upregulated significant (log2FoldChange > 0.25; Wilcoxon Rank Sum Test, Bonferroni-adjusted p-value < 0.05) differentially expressed genes (DEGs) enriched in pancreas ECs against other tissue ECs. The x-axis is average log2FC of the gene and the y-axis is –log10 of adjusted p-value. Top 5 DEGs for each plot are highlighted. B) Demonstrates overlap of all pancreas-enriched significant DEGs derived from the Tabula Sapiens EC atlas and a list of endothelial-enriched DEGs curated for this study from four different pancreas datasets across different age and species. This demonstrates pancreas-enriched DEGs pancreas-specific EC signature genes are in blue; other colors represent different overlaps of endothelial enriched and pancreas-enriched DEGs between the Tabula Sapiens, Descartes and Tabula Muris gene expression atlases. C) Demonstrates the overlap of all pancreas-enriched DEGs obtained from the Tabula Sapiens EC atlas with the curated list of endothelial-enriched DEGs. D) represents different overlaps of endothelial-specific pancreas-enriched DEGs between the Tabula Sapiens, Descartes and Tabula Muris EC gene expression atlases.

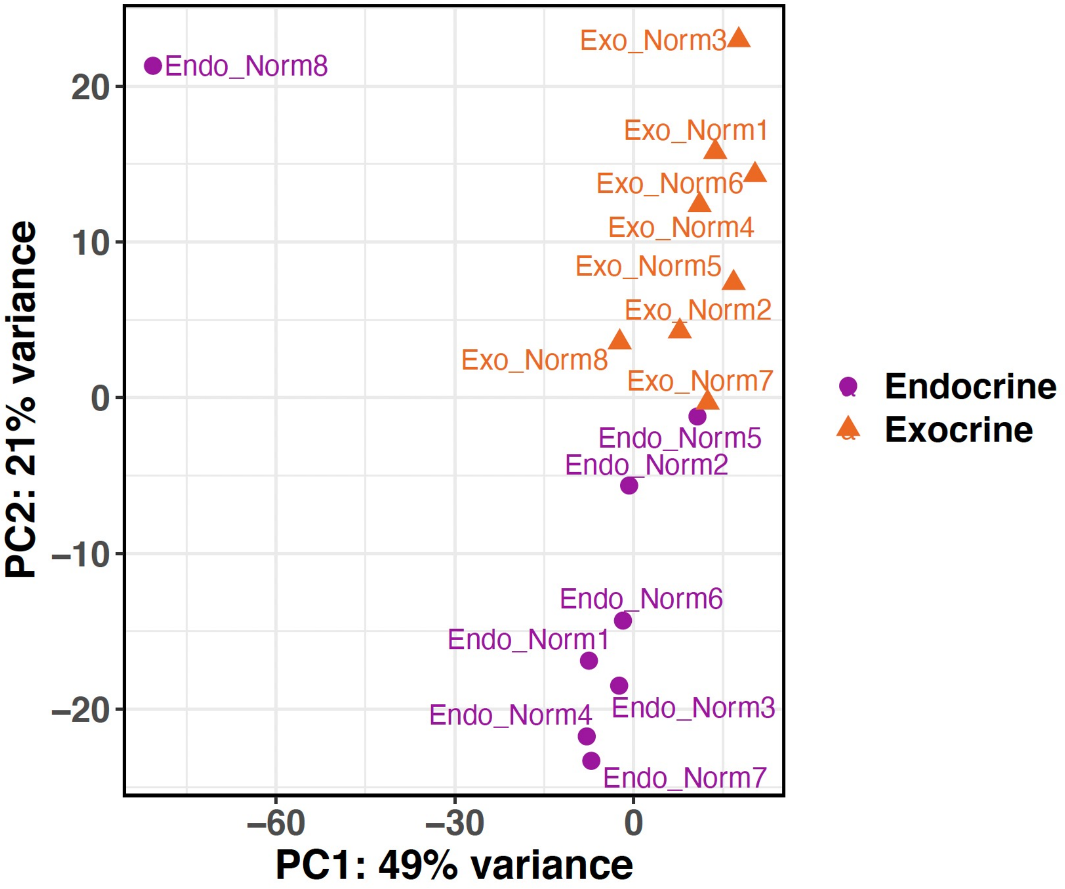


**Supplementary Figure 6. Principal Component Analysis of Bulk RNA-seq data**. Principal component analysis (PCA) of exocrine and endocrine EC samples from 8 normoglycemic donors (Norm) bulk RNA sequenced by Jonsson et al.

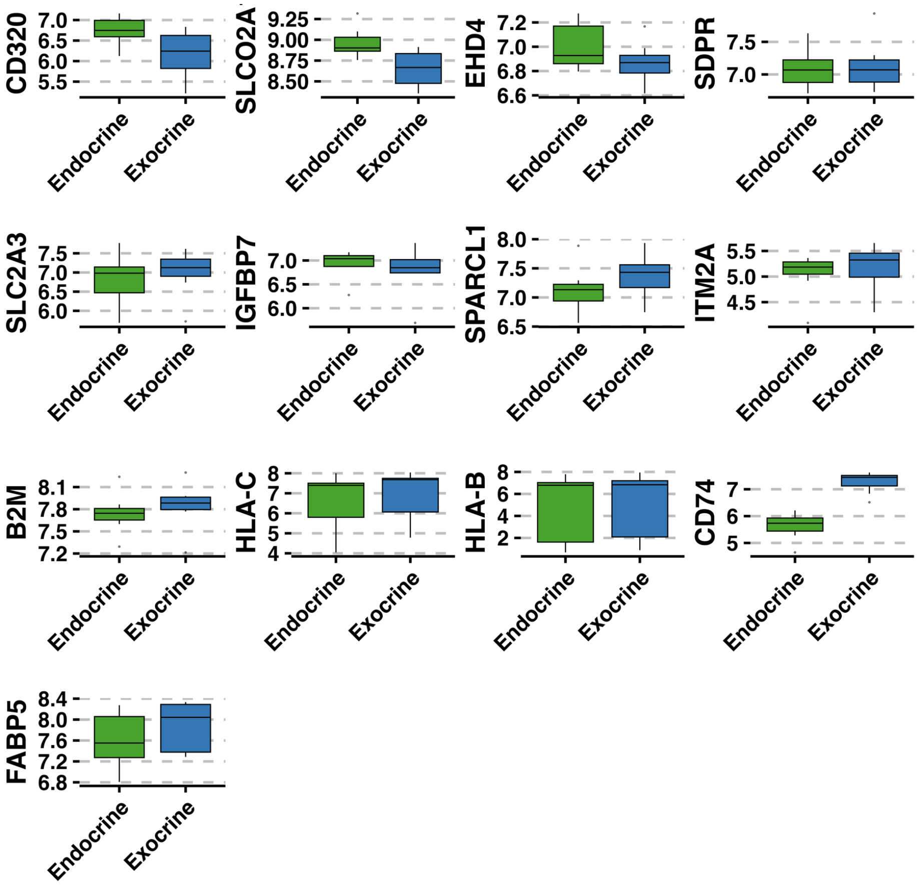


**Supplementary Figure 7. Assessment of non-significant genes of interest in bulk RNA-seq data**. Genes of interest enriched in capillary 1 compared to capillary 2 in the integrated pancreatic EC atlas that were not significantly (adjusted p-value > 0.05) enriched in endocrine EC samples compared to exocrine EC samples from the bulk RNA-seq data sequenced by Jonsonn et al. are shown

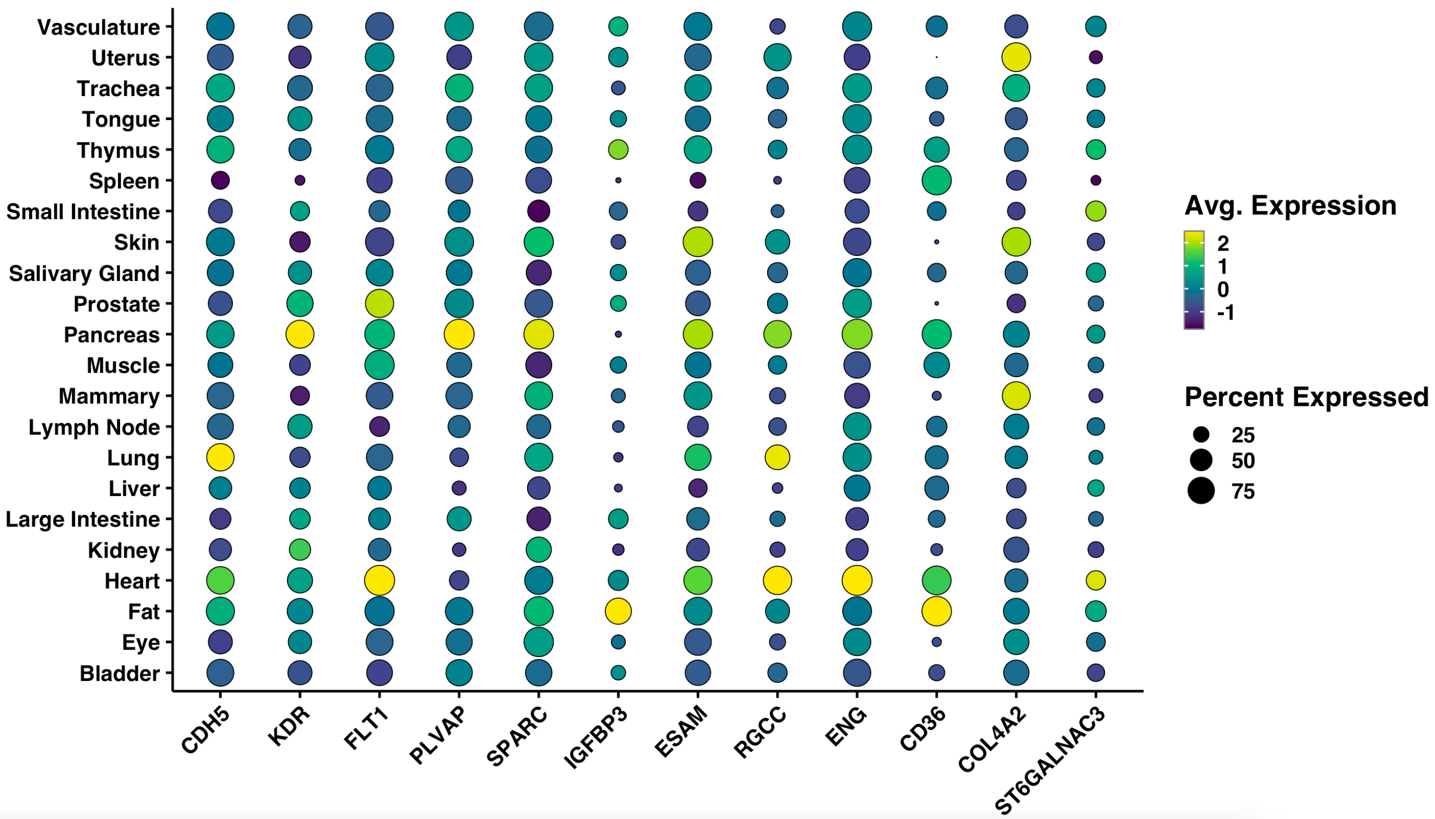


**Supplementary Figure 8. Assessing pancreas-specificity of markers identified for islet ECs in the literature**. Islet EC markers identified in the literature, where authors conducting differential expression analysis of endothelial population against other cell-types in the islet are shown in the Tabula Sapiens EC atlas.
